# Common AAV gene therapy vectors show indiscriminate transduction of living human brain cell types

**DOI:** 10.1101/2024.11.14.623624

**Authors:** JP McGinnis, Joshua Ortiz-Guzman, Maria Camila Guevara, Sai Mallannagari, Benjamin D. W. Belfort, Suyang Bao, Snigdha Srivastava, Maria Morkas, Emily Ji, Angela Addison, Evelyne K. Tantry, Sarah Chen, Ying Wang, Zihong Chen, Kalman A. Katlowitz, Jeffrey J. Lange, Melissa M. Blessing, Carrie A. Mohila, M. Cecilia Ljungberg, Guillermo Aldave, Ali Jalali, Akash Patel, Sameer A. Sheth, Howard L. Weiner, Shankar Gopinath, Ganesh Rao, Akdes Serin Harmanci, Daniel Curry, Benjamin R. Arenkiel

**Affiliations:** Department of Neurosurgery, Baylor College of Medicine, Houston, TX 77030, USA; Department of Molecular and Human Genetics, Baylor College of Medicine, Houston, TX, 77030, USA; Jan and Dan Duncan Neurological Research Institute, Texas Children’s Hospital, Houston, TX, 77030, USA; Medical Scientist Training Program, Baylor College of Medicine, Houston, TX 77030, USA; Development, Disease Models & Therapeutics Graduate Program, Baylor College of Medicine, Houston, TX 77030, USA; Stowers Institute for Medical Research, Kansas City, MO 64110, USA; Department of Pathology, Texas Children’s Hospital, Baylor College of Medicine, Houston, TX 77030, USA; Department of Pediatrics, Texas Children’s Hospital, Baylor College of Medicine, Houston, TX 77030, USA; Department of Neurosurgery, Texas Children’s Hospital, Baylor College of Medicine, Houston, TX 77030, USA

## Abstract

The development of cell-type-specific gene therapy vectors for treating neurological diseases holds great promise, but has relied on animal models with limited translational utility. We have adapted an *ex vivo* organotypic model to evaluate adeno-associated virus (AAV) transduction properties in living slices of human brain tissue. Using fluorescent reporter expression and single-nucleus RNA sequencing, we found that common AAV vectors show broad transduction of normal cell types, with protein expression most apparent in astrocytes; this work introduces a pipeline for identifying and optimizing AAV gene therapy vectors in human brain samples.

## Main

There is growing recognition that the field of medicine will soon see widespread use of molecular tools, including CRISPR-based gene editing, optogenetics, and other DNA– and RNA-based therapies. These new therapies hold great promise for treating some of our most challenging neurological diseases, from movement disorders and neurodegenerative diseases to epilepsies and tumors (Patel, Neurotherapeutics, 2024; Kreatsoulas, J Neurooncol 2024). However, in many cases, the safety and efficacy of these tools will depend on our ability to deliver sequence-based therapeutics precisely to target cell types, while avoiding neighboring but uninvolved cells. Viral vectors, including adeno-associated viruses (AAVs)—the most common vectors in ongoing clinical trials for neurological diseases (Lonser, JNS, 2020)—rely on their surface proteins’ recognition of cellular membrane components to gain entry to a cell (Challis et al., Annual Reviews Neuroscience, 2022). For this reason, modifying AAV capsids in order to target specific cell types (that is, modifying an AAV’s ‘tropism’) has been an area of broad interest.

Past efforts to screen and develop new AAV capsids for cell-type specificity have relied on animal models or cell lines (e.g., Pekrun, JCI Insight 2019; Kumar et al., Nature Methods, 2020; Ogden et al., Science, 2019; Li and Samulski, Nature Reviews Genetics, 2020), and the limited post-mortem human data following intraparenchymal AAV injection trials have not focused on identifying the cell types transduced (Bartus et al, Movement Disorders, 2011; Chu et al., Brain 2020, Castle, Human Gene Therapy, 2020). We therefore sought to use living human brain tissue to 1) establish the cellular specificity of FDA-approved (AAV2 and AAV9) and other common AAV variants in human brain tissue, identifying any tropism biases that could serve as the basis for iterative approaches to optimization; 2) create a standardized baseline against which to compare future novel capsids; and 3) identify potential species-specific tropism differences that would inform future AAV capsid engineering.

Towards this, we have adapted a human brain organotypic slice model (Figure 1a) using resected neurosurgical specimens not needed for pathologic diagnosis, and maintained them in a living state for two weeks *in vitro* (Chaichana 2007, Park 2022; Ting JoVE 2018, Ting Scientific Reports 2018; Schwarz Scientific Reports 2017, Schwarz eLife 2019, Bak et al., J Neuroscience Methods 2024). We determined the tropism profiles of fourteen common AAV vectors via a combination of immunofluorescence and single-nucleus RNA sequencing. We used eight natural AAV variants (AAV1, 2, 5, 6, 7, 8, 9, rh10), and six engineered AAV variants (DJ8, DJ, AAV2-retro, PHP.S, PHP.eB, Sch9) (Supplemental Table 1), all carrying eGFP constructs driven by the ubiquitous CAG promoter (ITR-CAG-eGFP-WPRE-hGH polyA signal-ITR) (Figure 1b). We chose the CAG promoter given its prevalence in preclinical and clinical gene therapy studies, and its strong and ubiquituous expression pattern. We used brain tissue from eight patients that included temporal lobe cortex and other lobectomy specimens resected due to intractable epilepsies, as well as normal cortex resected en route to deeper tumors (Figure 1c). AAVs were transduced at a uniform titer (2.1E9 vg per tissue slice) established to approximate the viral dose seen by brain cells infused during convection-enhanced delivery trials (Supplemental Table 2). Reasoning that capsid entry (i.e., AAV DNA presence) is therapeutically useful only to the extent that it leads to productive transcription and translation, we focused on identifying cell types that showed reporter RNA and protein expression fourteen days after virus transduction (where virus transduction occurred on post-operative day 1).

**Figure 1.**
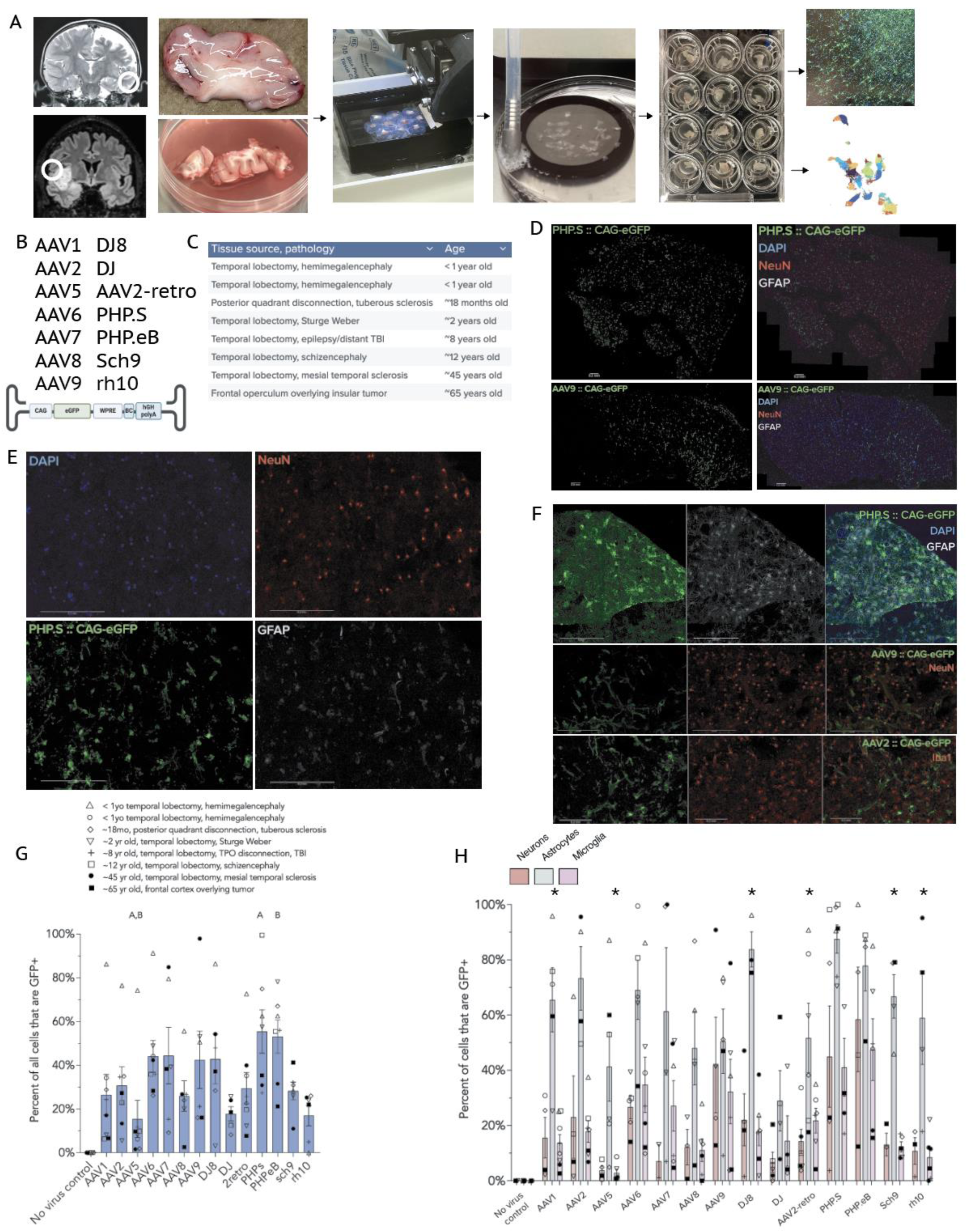
| Human brain organotypic slice model facilitates characterization of AAV tropism by immunofluorescence. **a**, Workflow of the human brain organotypic model: example MRIs with location of tissue resected, resected samples, vibratome slicing, NMDG-aCSF recovery, culture plates, and immunostaining or single-nucleus sequencing. **b,** The AAV capsid variants studied and the CAG-eGFP construct used for immunostaining and single-nucleus sequencing. **c**, The anatomical locations of patients’ samples, pathologies involved, and approximate ages. **d**, Representative images of entire sections of PHP.S and AAV9 from adult frontal cortex overlying tumor. Scale bars = 200 µm. **e,** Magnified images of PHP.S with DAPI, NeuN (neurons), GFAP (astrocytes), and transduced cells (GFP). **f**, Representative magnified images showing PHP.S and astrocytes, AAV9 and neurons, and AAV2 and microglia in pediatric temporal lobectomy samples. Scale bars = 200 µm. **g**, Percent transduction of all cells present in the tissue (measured as percent of DAPI-based ROIs that show GFP signal higher than the brightest control ROIs). (‘A’ indicates significant differences from the other bar labeled ‘A’, as does ‘B’. Kruskal-Wallis test with Dunn correction for multiple comparisons, adjusted p < 0.05) **h**, Percent of NeuN-positive, GFAP-positive, and Iba1-positive cells that co-express GFP, indicating successful transduction by the respective capsid variants. (* indicates p < 0.05 for post-hoc adjusted comparison of astrocytes versus microglia for the indicated variants except AAV2-retro, where the post-hoc significant comparison was astrocytes versus neurons. Kruskal-Wallis test with Dunn post-hoc test.)

For immunofluorescence analysis, the 300 µM cultured tissue slices were further cryostat-sectioned to 20 µM to facilitate high-resolution imaging. We systematically imaged the two 20 µM sections with the highest GFP expression levels (typically the most superficial sections). Since many slices showed GFP expression eccentrically, we restricted analysis to a 2 mm circle centered around the highest GFP expression for each image to standardize analysis (Supplemental Figure 1a,b) (see Methods for detail). These measures were taken with the rationale that we sought to identify each AAV’s transduction efficacy at their presumed point of highest concentration, and mitigate the variability that was introduced by differing slice sizes. While this does have the effect of increasing the apparent percent of cells transduced, it retains the relative relationships between the AAV variants at their point of greatest transduction.

Of the fourteen variants tested by immunofluorescence, PHP.eB and PHP.S (55% and 53%, *n* = 7 tissue samples per variant) showed the greatest overall transduction rates (measured as percent of DAPI-based ROIs that were also GFP^+^; Kruskal-Wallis test with Dunn post-hoc test) (Figure 1d, 1g). AAV5, DJ, and rh10 showed the lowest transduction rates, all averaging less than 20% of the total cells in the quantified area (*n* = 5-8). AAV2 and AAV9—the most common AAV variants used in neurological trials—both showed moderate transduction, around 30 and 40% of cells within the analyzed area, respectively (*n* = 7 and 6, respectively).

For general neuronal transduction (measured as percent of NeuN^+^ cells that also expressed GFP; numbers of tissue samples were lower due to some cases where the highest GFP^+^-containing area did not contain sufficient numbers of neurons for evaluation), PHP.eB and PHP.S showed the greatest transduction rate (∼55% and 45%, *n* = 4 and 5, respectively), with AAV9 and AAV2 transducing ∼40% and ∼30% of NeuN^+^ neurons respectively (*n* = 4 for each). AAV5 and DJ showed the lowest levels of neuronal transduction (∼5%, *n* = 4 and 5) (Figure 1h). These patterns are notably different from several studies of mouse cellular tropism (Aschauer, PLoS ONE 2013; He, Human Gene Therapy Clinical Development, 2019).

Astrocytes (GFAP^+^ cells) were notably the most highly transduced cell type for nearly all capsid variants tested. AAV1, AAV2, AAV6, AAV7, DJ8, PHP.S, PHP.eB, and sch9 all transduced ∼60% or more of astrocytes within the areas of analysis, while rh10 and AAV2-retro both transduced just more than 50% of astrocytes analyzed (Figure 1h, *n* = 3-8). The lowest rate of astrocyte transduction in the analyzed areas was observed with DJ and AAV5, ∼30-40% (*n* = 5-6).

Microglia (Iba1^+^ cells) showed variable transduction, from as low as ∼3-15% with AAV5, AAV8, and DJ, to as high as 45% with PHP.S and PHP.eB (Figure 1h, *n* = 5-6).

In general, variants that exhibited higher rates of transduction (e.g., PHP.eB, PHP.S) tended to show higher rates of transduction across all cell types, while variants with lower transduction rates (AAV5, DJ) tended to show lower transduction across all cell types. Most capsid variants, including the FDA-approved AAV2 and AAV9 serotypes, showed moderate transduction, averaging 30-50% of all cells transduced. Finally, astrocytes tended to show higher rates of transduction than either neurons or microglia.

Because future AAV screens using human brain tissue will require high throughput methods capable of accommodating libraries of hundreds of thousands of AAV variants, we next sought to determine whether single-nucleus RNA sequencing could capture barcoded versions of these same fourteen AAV variants from two patient samples (Supplemental Table 1). The first sample was temporal lobe cortex from a pediatric patient with Sturge-Weber syndrome (a neurocutaneous syndrome frequently causing epilepsy), while the second sample was adult frontal opercular cortex overlying an insular glioma (Figure 1a). Each tissue was sectioned to 300 µM slices; six slices per patient were dosed each with a 4 µL droplet containing the combined library at a total of 2.1E9 vector genomes (each variant dosed at 2.1E9 * 1/14, though the library used for the pediatric temporal lobectomy sample was missing sch9 and therefore each remaining virus was 2.1E9 * 1/13). Six control slices were dosed with a 4 µL drop of PBS. All samples again were kept in culture until fourteen days after virus application, then processed for single-nucleus RNA sequencing.

To assess whether cultured tissue remained reasonably representative of human brain cell types over the two weeks, we generated combined and subsetted UMAP plots that included all cells from day 0, day 14 with virus, and day 14 without virus. We noted similar clustering between control day 0 samples and day 14 samples both with and without virus, albeit with some loss of neurons at day 14 (Figure 2a, Supplemental Figure 2a). Assessing the global transcriptional pattern of each cell type across all significantly variable genes, we found reasonable correlations between the log2 fold change for the day 0 and day 14 transduced samples (Figure 2b), indicating that the distinctive identities of each cell type were reasonably well preserved across two weeks in culture.

**Figure 2.**
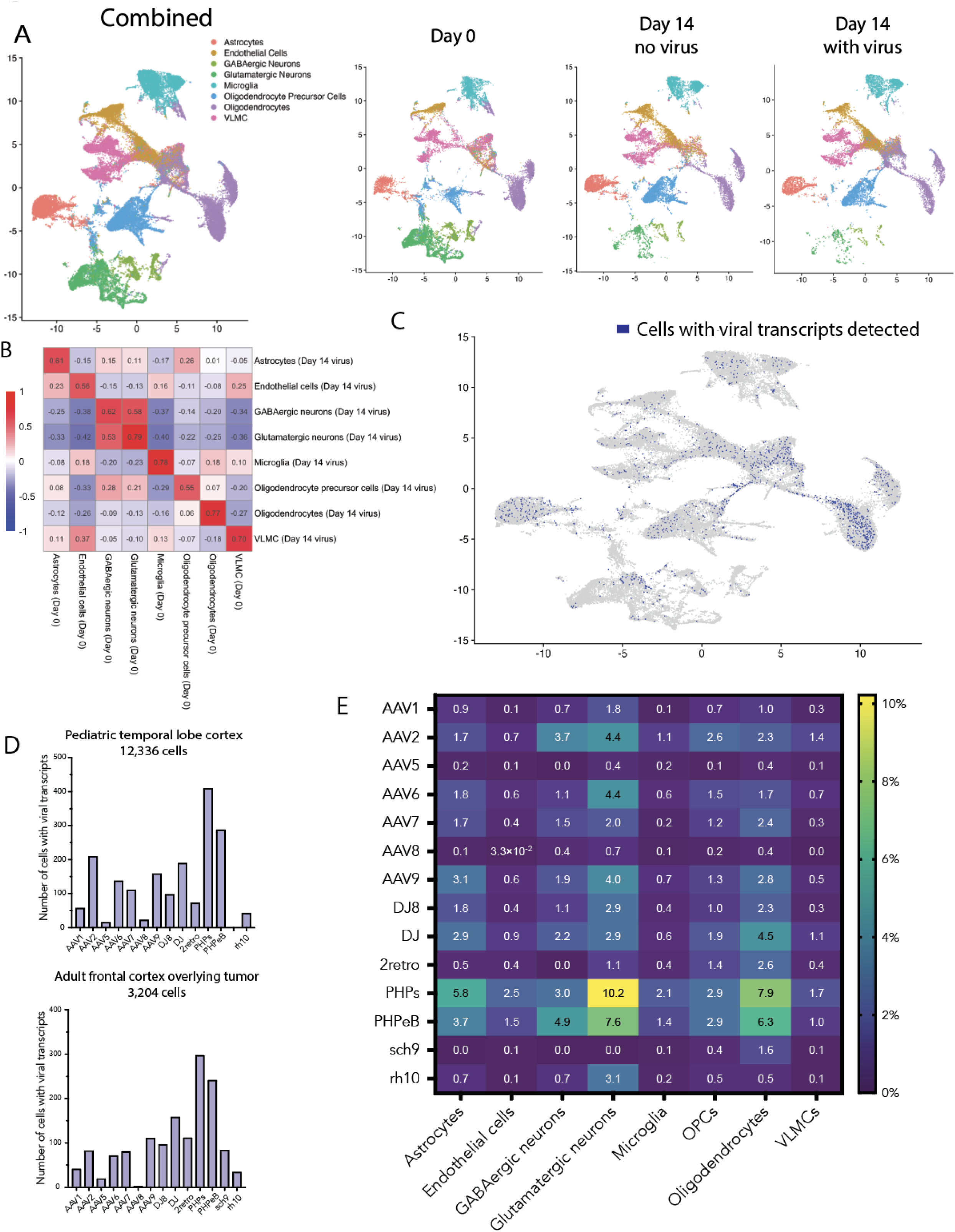
| Single-nucleus RNA sequencing identifies human brain cell types transduced with AAV capsid variants. **a**, Samples from the pediatric temporal lobe and adult frontal lobe were sequenced and aligned independently and then combined for cluster analysis. Panels separated by day. Clusters assigned using SingleR and the Allen Institute’s middle temporal gyrus transcriptomic data. **b,** Correlation matrix for log2 fold change by cell type, comparing day 0 tissue to day 14 transduced (virus added) tissue. Each cell contains the Pearson correlation coefficient. **c,** UMAP of the Day 14 with virus sample (15,540 cells), with dark blue cells indicating the presence of at least one viral transcript identified (1,432 cells). **d,** Bar graphs showing total number of cells transduced by each AAV capsid variant for the pediatric and adult samples (pediatric sample missing Sch9). **e,** Heatmap showing percent of each cell type transduced by AAV capsid variant.

A total of 1,887 cells contained at least one viral transcript (out of 15,540 total cells) (Figure 2c). We noted good concordance of the relative transduction efficiency between the best and worst AAV variants in both the pediatric and adult single-nucleus RNA sequencing samples, as well as with our imaging data (Figure 2d compared to ‘all cells’ [DAPI] bars in Figure 1g). Plotting separate UMAPs for each variant revealed transduction of most major cell clusters by each variant, again indicating a lack of cell-type-specific tropism for the tested AAV variants (Supplemental Figure 3). We next calculated the percent transduction for each cell type by variant (Figure 2e). Given the reduced titer of each variant in the combined library (1/14^th^ of the dose used for immunofluorescence), the use of entire tissue sections for sequencing instead of focused analysis (as for imaging), sequencing depth of ∼50,000 reads per cell, and dropout effects, our absolute transduction levels seen by sequencing were expected to be significantly lower than our imaging results—however, we note that absolute levels here were of less concern than the relative transduction efficiency between variants and across cell types.

Overall, we found, using ex vivo living human brain tissue, that these fourteen common AAV gene therapy vectors show largely indiscriminate transduction across cell types, though transduction efficiency varies considerably by AAV capsid variant. PHP.eB and PHP.S showed the greatest transduction rates both by imaging and by single-nucleus RNA sequencing, while AAV5 was consistently among the least efficient AAVs. Notably, while PHP.eB was recognized for its broad transduction in the mouse brain, PHP.S showed (in mice) minimal brain transduction apart from the brain stem, and was instead recognized for efficient targeting of sensory neurons, indicating species-specific tropism differences when compared with our human data here (Chan et al., Nature Neuroscience, 2017). The two variants in clinical use, AAV2 and AAV9, showed moderate transduction by imaging and single-nucleus RNA sequencing.

Transgene protein expression (GFP) was generally most prominent in astrocytes, though transgene mRNA seems more evenly distributed. In order to promote strong protein expression for improved gene therapies, the balance of factors that control transgene expression in human brain cell types will need to be a focus of future studies (Schwaunhausser et al., Nature, 2011, Gonzalez-Sandoval et al., Nature Comm, 2023, Loeb et al., Cell Reports, 2024). We note that the indiscriminate transduction of these AAV capsid variants may be due to their initial identification in mice (i.e., the variants may share features that helped them succeed in those screens), which in most cases have been selected for broad CNS transduction; moreover, several are closely related.

We have only used a single promoter, CAG, one of the most common in neurological clinical trials, as well as in pre-clinical animal work. The rationale for this is that when assessing capsids for their cell type specificity, one should use the broadest and strongest promoter (and in this case one of the most clinically relevant promoters); we wanted to examine the cell type specificity of the capsids in this study, not a combination of the capsid and promoter. However, it will certainly be the case that various promoters, in combination with specific AAV capsid variants, will help narrow cell-type-specific expression. Promoter identification and expression analysis in human brain tissue is therefore the focus of ongoing work.

With respect to the translational value of this study, the lack of pial or ependymal layers covering our slices means that this method most closely approximates direct intraparenchymal injections rather than systemic or intraventricular delivery, and should be interpreted accordingly (Harkins, Neurotherapeutics, 2024). Our findings indicate that intraparenchymal gene therapy injections with these capsids can be expected, when carrying the CAG promoter, to transduce most cell types present, with notably strong protein expression in astrocytes.

Finally, human patient samples are reasonably expected to be more heterogeneous than laboratory strains of inbred mice or cell culture lines, and so more variability should be expected, which may be considered a limitation. As others have argued, however, testing therapies in these more variable but relevant conditions should help identify interventions whose signal is able to surmount such variability, and therefore predict clinical trial performance more reliably (Park et al., 2022; Scannell and Bosley, PLoS ONE, 2016). Together, these findings underscore the continued need to develop cell-type-specific AAV variants, and the importance of using the ultimate intended target of such vectors, human brain tissue, for precise tropism characterization (Pupo et al., Molecular Therapy, 2022; Zhu et al., Scientific Advances, 2024).

## Methods

### Human brain tissue samples

Patients planned to undergo resective neurosurgical procedures at three major neighboring academic hospitals were approached and consented pre-operatively by a member of the study team (Baylor College of Medicine IRB H-51865) for any specimens not needed for pathologic diagnosis. In the case of pediatric patients or adults unable to consent, parents or medical decision-makers were consented.

Specimens were collected from pathology following tissue sectioning and gross examination by board-certified neuropathologists, or directly from the operating room in sterile plastic specimen containers, usually within minutes and no longer than 45 minutes after resection, into which we poured ice-cold, pre-carbogenated (>20 min bubbling time, 95%/5%) NMDG-aCSF, following the protocol from Park et al. (Park, Nature Protocols 2022). Tissue was rapidly transported to the lab on ice, where it was manually sectioned with scalpels into pieces ∼1cm x 1cm. Tissue pieces were placed into a retractable tube and molten ultra-low melting agarose (Sigma A2576) was poured over top. Using a cold clamp the agarose was rapidly solidified, at which point the agarose cylinder was removed and superglued onto a tape-covered vibratome mount. Tissues were oriented so that the sectioning would be done perpendicular to the cortical surface, parallel to neuronal projections. Tissues were sectioned to 300um on a Leica VT1200 vibratome in cold aCSF that was being continuously bubbled with carbogen. We then moved the slices to room temperature aCSF that was continuously bubbled with carbogen for 15-30 minutes, after which the slices were plated individually onto membranes inserts(CellTreat #230617) overlying 600 µL of organotypic slice culture media (OSCM) as detailed in Park et al., 2022. Any excess media or aCSF was aspirated off the membrane surface, and the plates were kept in a humidified tissue culture incubator at 37 °C, 95% humidity, and 5% CO2. The tissue underwent daily media changes (400 µL removed and added to the surrounding well, avoiding any contact with the tissue slice).

### AAV Production

All AAVs were packaged in-house by the Texas Children’s Hospital Jan and Dan Duncan Neurological Research Institute’s Neuroconnectivity Core (supported via NIH P50HD103555). Viral capsid gene sequences are found in Supplemental Table 1. The CAG-EGFP-WPRE-hGH polyA signal was modified by removing the bGH polyA signal from Addgene #37825 (gift from Ed Boyden) and replacing it with the hGH polyA signal. The entire plasmid sequence is found in Supplemental Table 1.

### Cell Culture and Transfection

A three-vector system was used for AAV production (Cell Biolabs). HEK-293 cells were plated in 15 cm dishes at a density that yielded ∼70% confluency the following day. Cells were then transfected in each plate with 25μg helper plasmid, 25 μg serotype specific AAV vector, and 25 μg of AAV shuttle vector using polyethylenimine (PEI). In a 1:3 ratio (µg DNA:µg PEI), the solution was added dropwise to cells. After 4-6 hours, the medium was changed to DMEM, 5% FBS, 1x Penicillin/Streptomycin. 48-72 hours later, transfected cells were harvested using a cell scraper. Cells were pelleted by centrifugation at 3500rpm for 10 minutes at 4°C. The supernatant was removed, and the pellet resuspended in TMN (50 mM Tris pH 8.0, 5 mM MgCl_2_, 0.15M NaCl) at a concentration of 1ml/plate. The resuspended cells were frozen at –80°C overnight.

### Purification

10 µL of DNaseI (10 mg/ml) and 10 µL RNase A (1mg/ml) were added to each plate of defrosted cells in TMN. Plates were incubated at 37 °C for 30 minutes, shaking frequently. 100 μl of 5% sodium deoxycholate solution in water was added to each plate and mixed gently. Plates were then incubated at 37 °C for 10 minutes. The suspended samples were transferred from the plates to tubes and placed on ice for 15 minutes. The tubes were then centrifuged at 3700rpm for 10 minutes and the supernatant was collected.

### Iodixanol Gradient

OptiPrep™ (Millipore Sigma D1556-250ML), or iodixanol, was purchased as a 60% (W/V) stock in water. 15%, 25%, and 40% dilutions of iodixanol were made in PBS-MK (1x PBS, 1 mM MgCl_2_, 2.5 mM KCl). 2.5μl phenol red solution (0.5% stock in PBS-MK) was added per 1ml of iodixanol solution in the 25% and 60% fractions. The gradient was loaded to the bottom of Beckman OptiSeal 16X67mm tubes (Cat# 362181) starting with 1.5ml 15% iodixanol, 1.3 ml 25%, 1.4ml 40% and finally 1.3ml 60% iodixanol. The supernatant collected from the previous purification step was then placed on top of the gradient. Tubes were centrifuged at 60000rpm for 90 minutes in a Beckman NVT 65 rotor. The clear band below the 60% mark (and below the white cellular debris layer) was collected using a needle and syringe. The collected volume from each tube was approximately 1.5ml.

### Concentration

The goal of this step was to remove the OptiPrep and concentrate the AAV using an Amicon Ultra-15 Centrifugal Filter (Millipore Sigma, UFC9 100 24). An Amicon column was equilibrated with 15ml of DPBS (no Mg, no Ca) by centrifugation at 2500rpm for 5 minutes. The collected band from the OptiPrep gradient was mixed with approximately 40ml of DPBS. The samples were run in batches through the Amicon filter, discarding the filtrate between spins. The virus was then washed three times with 15ml of DPBS with centrifugation after each wash at 2500rpm for 10 minutes. The virus was collected to a sterile microcentrifuge tube, aliquoted, and frozen to –80°C.

### Viral Titer

Titering of virus was performed using Applied Biological Materials qPCR AAV Titer Kit (Cat# G931) and following the manufacturer’s recommended protocol. Viral preparations were first diluted to ∼10^8^ GC/mL before undergoing viral lysis at room temperature for 3 minutes. A standard curve was generated using five 10-fold serial dilutions of provided Standard Control DNA (dilutions 1/100 to 1/100,000). qPCR components and cycling conditions are found within the manufacturer’s accompanying product datasheet. Final titer analysis was performed using the manufacturer’s provided calculation file.

### Virus application

On the day after collection (post-operative day 1), 2.1E9 virus is applied (in a volume of 4 µL where sterile PBS is used to supplement volume up to 4 µL) to the surface of the slice. The 4µL droplet at the end of a 10 µL pipette tip is touched to the center of the slice. The media underlying the slices underwent daily changes until fourteen days after virus addition, at which time they were either flash frozen in liquid nitrogen (for later single-nucleus isolation and sequencing) or fixed in 4% PFA in PBS (for imaging). Control slices had 4 µL of PBS pipetted instead of AAVs. For imaging, one variant per well was used, at a dose of 2.1E9. For the combined pool for sequencing, the total dose was 2.1E9, such that each variant was present at 1/14 * 2.1E9 for the adult sample and 1/13 * 2.1E9 for the pediatric sample (which was missing Sch9).

### Histology, imaging, and analysis

Fourteen days after virus addition, tissues were fixed in 4% PFA overnight at 4 °C by removing media from the wells and adding 4% PFA; the next day, PFA was removed and replaced with 20% sucrose overnight and then 30% sucrose overnight for cryopreservation. Tissues were then mounted exposed surface down in OCT such that the first 20 µM cryostat slices would be the surface on which virus was applied. Sections were affixed to typically 4 slides sequentially, such that each slide contained some superficial, middle, and deeper layers of each tissue slice. Slides were stored in –20. Slides were rinsed with PBS-0.1% Triton-X for ten minutes, then blocked overnight at 4 °C in PBS-0.1%T+10% normal donkey serum. Slides were then placed tissue-side down in thin slide folders (Amazon OakRidge Products B00X6L1NM4) over 600uL of combined 1:1000 chicken anti-GFP (Abcam 13970), rabbit anti-NeuN (Abcam 177487), and mouse anti-GFAP (Abcam 279290) or 1:500 rabbit anti-Iba1 (ThermoFisher PA5-27436) overnight at 4 °C. Slides were washed five times for at least five minutes each time in PBS-0.1%T, and similarly placed on 600 µL secondary antibody (1:1000 goat anti-chicken Alexa 488, goat anti-rabbit Alexa 546, and goat anti-mouse Alexa 633). Slides were again washed 5x for at least 5 minutes each, partially dried, covered with fluoromount G with DAPI, and coverslipped.

Approximately 4 to 6 sections were present per slide depending on section size. All sections were inspected visually under 10X GFP epifluorescence to confirm similarity, and the two sections with highest GFP intensity were selected for complete confocal imaging on a Leica SP8 microscope using a 10X objective/NA 0.4. Because we care most about approximating, as best as possible, the percent of cell type transduction at the tip of a catheter used for intraparenchymal injection, we wanted to analyze the best possible transduction capabilities of each virus. We also needed to control for variability in slice size and eccentric transduction due to liquid slide/spread of the pipette drop of virus. In order to mitigate these factors, a consistent 2mm circular ROI was applied to every section and manually centered over the area of most dense GFP expression. (Example images are found in Supplemental Figure 1a with overlying ROIs that were used.) In this way, larger slices but with a similar area of transduction would not be penalized simply for being larger and having more untransduced (or more weakly transduced) areas—the brightest area of transduction on each slice would be analyzed similarly. This relies on the assumption that the area of brightest GFP signal is the area that saw the greatest amount of virus, which, while not certain, is a reasonable assumption.

Using a custom FIJI/ImageJ program (https://github.com/jpmcginnis/HumanAAVProject/blob/main/imageJ-imagesorterforlifloop), .lif files were converted to .tif files, the .tif files were automatically scanned by a standardized 2 mm circle to find the location that would maximize GFP signal within the circle (https://github.com/jpmcginnis/HumanAAVProject/blob/main/imageJ-automatedROIloopandclose). The program saved a screenshot of where that circle was placed, and then saved separately an image only of the data within the 2 mm circle. Using that restricted image, ROIs were drawn around each DAPI nucleus using another custom imageJ/FIJI program, and the average intensity across each individual ROI was calculated separately for each of the four channels. (Code: https://github.com/jpmcginnis/HumanAAVProject/blob/main/imageJ-Macro2-DAPIchannel1-loop.)

Thresholds for positive/negative GFP cells were assigned by identifying the maximally brightest ROI of the untransduced control slices for each separate patient sample, and setting the threshold at that value for that patient’s imaging analysis (such that any ROI brighter than the maximally bright ‘untransduced’ ROI would be called GFP^+^); the threshold was different for different patient samples (https://github.com/jpmcginnis/HumanAAVProject/blob/main/googlecolab-Step1-Ch1DAPICh2-633-Ch3-488-Ch4-546-generatepercentpositiveneedtosetthresholds). Thresholds for for NeuN^+^, GFAP^+^, and Iba1^+^ were set by examining representative images’ ROIs within that channel and selecting a threshold that would include clearly positive cells but disregard negative cells; this was typically around the mean value for the channel. Using custom Python language in Google Colabs, the ROIs were then identified as either positive or negative for GFP or the other channels, and the percent of a certain channel (DAPI, anti-rabbit 546, or anti-mouse 633) positive for GFP was calculated. In order for an image to be included, it needed to have at least 30 cells of that type to prevent conclusions being made from low numbers of cells. Graphs were created using Graphpad Prism 10. All code available at https://github.com/jpmcginnis/HumanAAVProject/tree/main.

### Single-nucleus RNA sequencing

For the tissue sections that received barcoded viral libraries (total titer 2.1E9 in 4 µL PBS per tissue slice), 14 days after virus application the tissue was gently removed from its well, flash frozen in liquid nitrogen, and stored at –80 °C. On the day of processing, frozen tissue was dissociated using GentleMACS nuclei isolation protocol (nuclei isolation buffer [Miltenyi Biotec, cat# 130-128-024], Protector RNAse Inhibitor [Millipore Sigma, cat# 3335402001], GentleMACS C tubes [Miltenyi Biotec, cat# 130-093-237], GentleMACS Octo Dissociator [Miltenyi Biotec, cat# 130-096-427], MACS SmartStrainers 70um [Miltenyi Biotec, cat# 130-098-462], MACS SmartStrainers 30um [Miltenyi Biotec, cat# 130-098-458]). In brief, samples were placed in 2 mL of Miltenyi Nuclei Isolation Buffer and Protector RNAse Inhibitor in GentleMACS C tubes. Samples then underwent the preprogrammed “nuclei isolation” program on a GentleMACS Octo Dissociator. Immediately after dissociation, samples were strained through a 70um MACS SmartStrainer and collected in a 15 ml tube, centrifuged at 300xg for 5 minutes at 4°C. The supernatant was extracted and discarded, and the resulting pellet resuspended in 1mL of ice-cold PBS. Resuspended samples were then run through a 30um MACS SmartStrainer, centrifuged, and resuspended in ∼250-500 µL. Nuclei were then FACS sorted using DAPI on a Sony MA900 in the Baylor Cytometry and Cell Sorting Core.

### Library Preparation

The single-cell gene expression libraries were prepared according to the Chromium Single Cell Gene Expression 3’v3.1 instruction (PN-1000121, PN-1000120, PN-1000213, 10x Genomics). Briefly, single cells, reverse transcription (RT) reagents, Gel Beads containing barcoded oligonucleotides, and oil were loaded on a Chromium controller (10x Genomics) to generate single-cell GEMS (Gel Beads-In-Emulsions) where full-length cDNA was synthesized and barcoded for each molecule of mRNA (UMI) and each single cell (cell barcode). Subsequently, the GEMS were broken and cDNA from each single cell was pooled. Following cleanup using Dynabeads MyOne Silane Beads, cDNA is amplified by PCR. The amplified product was fragmented to optimal size before end-repair, A-tailing, and adaptor ligation. The final library was generated by amplification.

### Cell Type Annotation and Barcode Identification

Fastq files were uploaded to the 10X Cloude Analysis web browser. A custom reference genome was created using human GRCh38 to which the fourteen 25 base pair barcode sequences were added (barcode sequences found in Supplemental Table 1); this was used to align the fastq files. R/Seurat: Each sample’s (day 0, day 14 without virus, day 14 with virus) filtered matrix HD5 file was downloaded from the 10X cloud and was used to generate a Seurat object. These Seurat objects were then integrated to create a combined object for joint UMAP creation. The Allen Institute Brain Atlas transcriptomics explorer cell type CSV for human middle temporal gyrus (<u>Gene expression by cell type, medians</u>) was downloaded and SingleR was used to assign cell types. These were then collapsed into general categories (Astrocyte, Glutamatergic neuron, Oligodendrocyte, etc.), and the FindMarkers() function in Seurat was used to identify differentially expressed genes between each cluster for a given sample by day (separately for day 0, day 14 without virus, and day 14 with virus). We generated a list of all genes showing differential expression between any two clusters for that given sample, log2fold changes, and p values, which was downloaded for each cell type, and then Pearson correlation was calculated across all differentially expressed genes between pairs of cell types; genes not present in one of the two cell types being compared were discarded (code at https://github.com/jpmcginnis/HumanAAVProject/tree/main). Graphs and UMAPs were created in Graphpad Prism 10 or using ggplot2.

### Statistical analysis

Given the lack of normality, imaging data was analyzed by nonparametric Kruskal-Wallis tests with Dunn post-hoc tests, adjusted for multiple comparisons where appropriate. *n* = 3 to 8 depending on the AAV variant and cell type because not all cell types were sufficiently represented (we required at least 30 of that cell type to be present to analyze that cell type for that tissue/variant). All statistical tests were two-sided with α=0.05. Data are presented as mean ± SEM unless otherwise indicated. For single-nucleus RNA sequencing analysis, differential expression analysis was performed using Seurat/FindMarkers() using the nonparametic Wilcoxon rank sum test with Benjamini-Hochberg correction for multiple comparisons.

## Funding

The majority of the funding for this project came from a generous gift from Mr. Michael Wilsey and family. The project was also supported in part by the NRI Neuroconnectivity and Viral Core (supported via NIH P50HD103555) and the RNA In Situ Hybridization Core facility at Baylor College of Medicine (supported by a Shared Instrumentation grant from the NIH 1S10OD016167 and the NIH IDDRC grant P50 HD103555 from the Eunice Kennedy Shriver National Institute Of Child Health & Human Development). This project was further supported by the Cytometry and Cell Sorting Core at Baylor College of Medicine with funding from the CPRIT Core Facility Support Award (CPRIT-RP180672), the NIH (CA125123 and RR024574) and the assistance of Joel M. Sederstrom, and by the Single Cell Genomics Core at Baylor College of Medicine with funding from the CPRIT RP200504 and the NIH S10OD032189.

## Acknowledgements

We thank the hundreds of patients who have generously participated in this project, as well as Ying, Rong, Hilary, Gemma, Megan, Jennifer, Loretta, Glenn, Omar, Waseem, Johannah, Neal, Mira, the Arenkiel lab, cytometry core, and the unnamed operating room, pathology, and research staff who have contributed so much time and effort to this work.

---

**Supplemental Figure 1.**
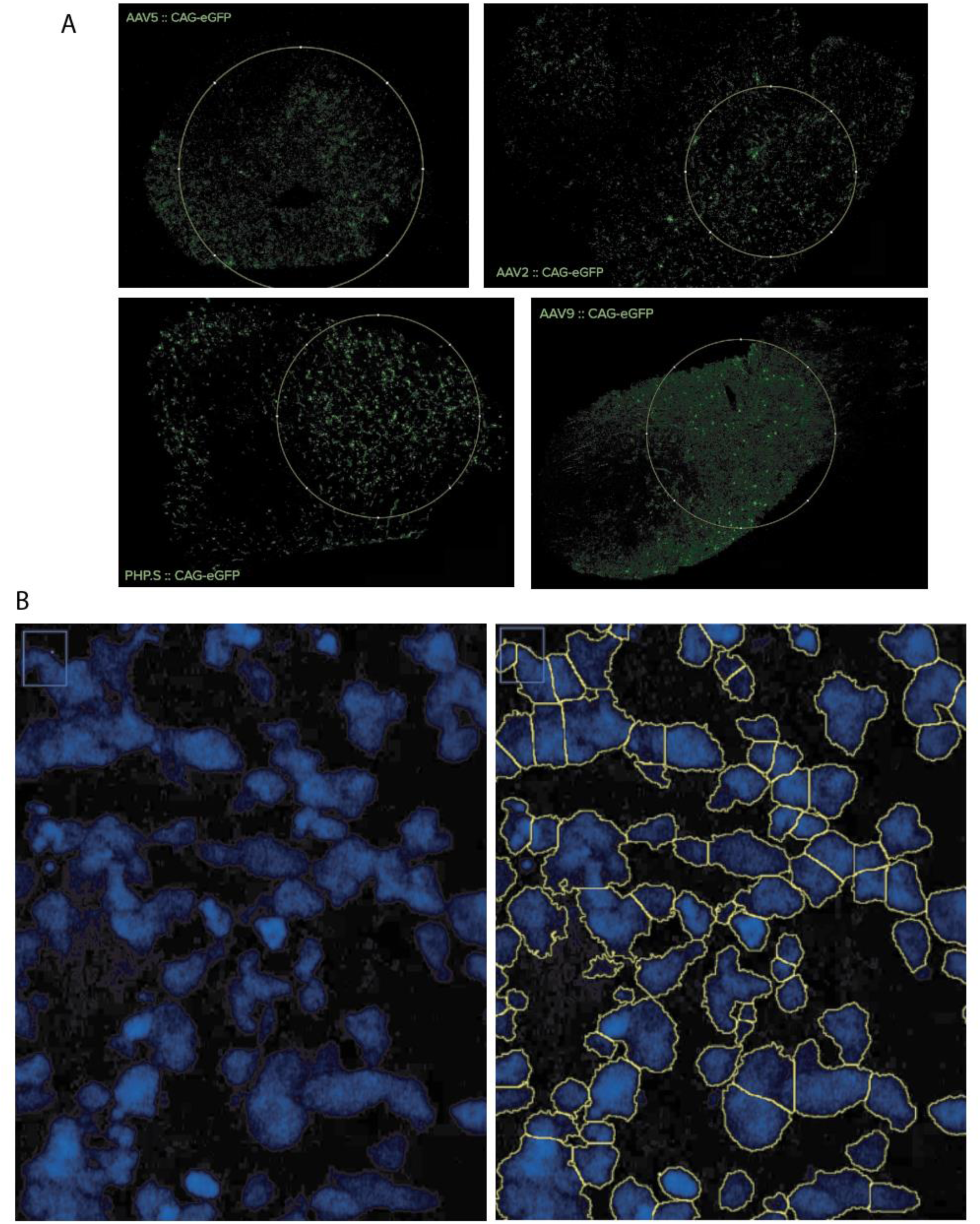
| Selection of regions of interest (ROI) for image analysis. **a**, Example screenshots of the 2mm circular region used to restrict analysis to the highest 2mm diameter circlular region of each slice. **b,** Representative images showing the custom automated DAPI-centric ROI identification. The average intensity within each ROI is separately calculated for each of the four channels.

**Supplemental Figure 2.**
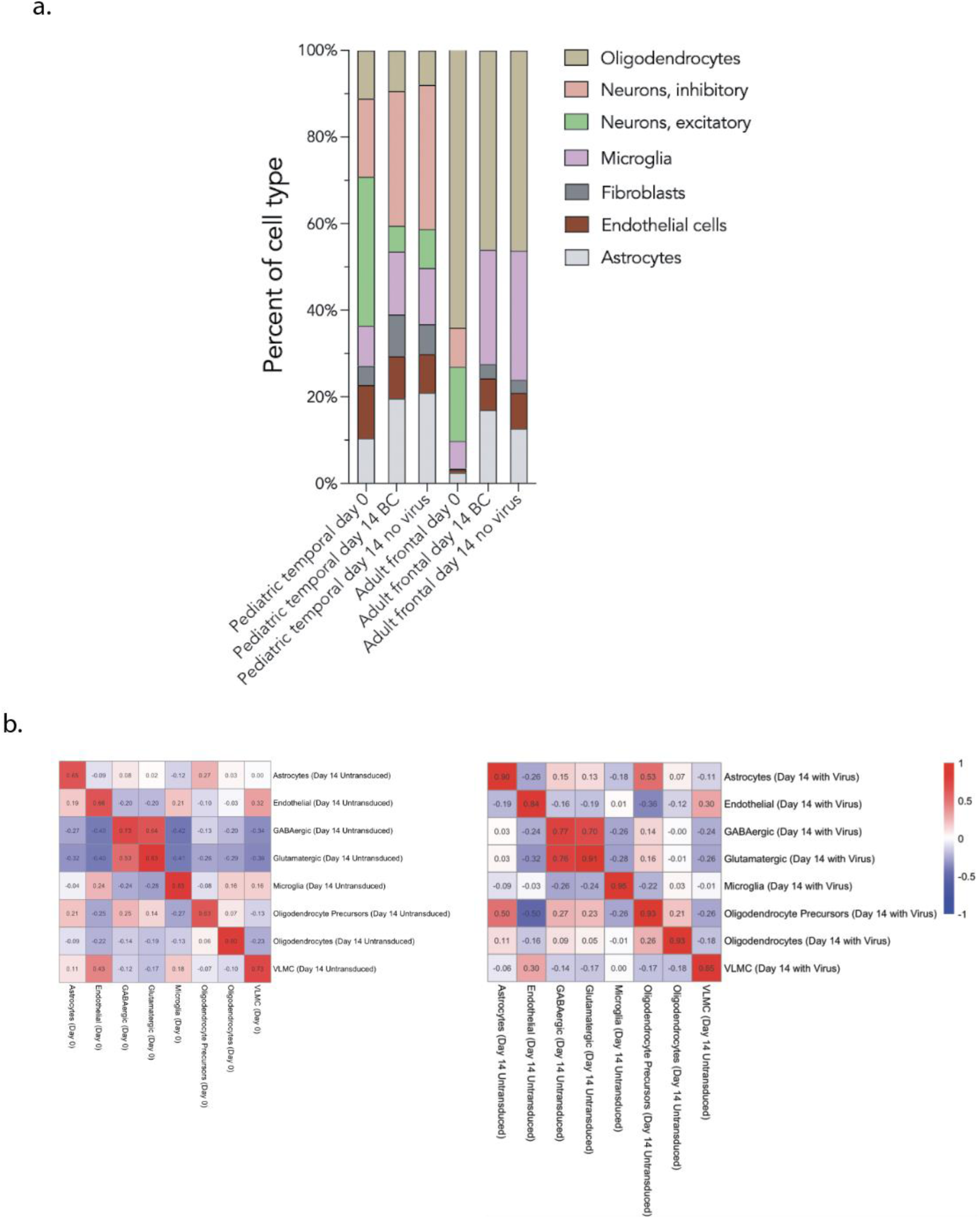
| Cell type composition changes across fourteen days. **a**, Proportions of cell types present in sequenced tissues at each time point, with and without virus. **b**, Correlation matrices (correlation calculated across all differentially expressed genes identified for that sample’s clusters) for day 0 versus day 14 without virus, and for day 14 with virus versus day 14 without virus. Pearson correlation coefficients are printed within each cell.

**Supplemental Figure 3.**
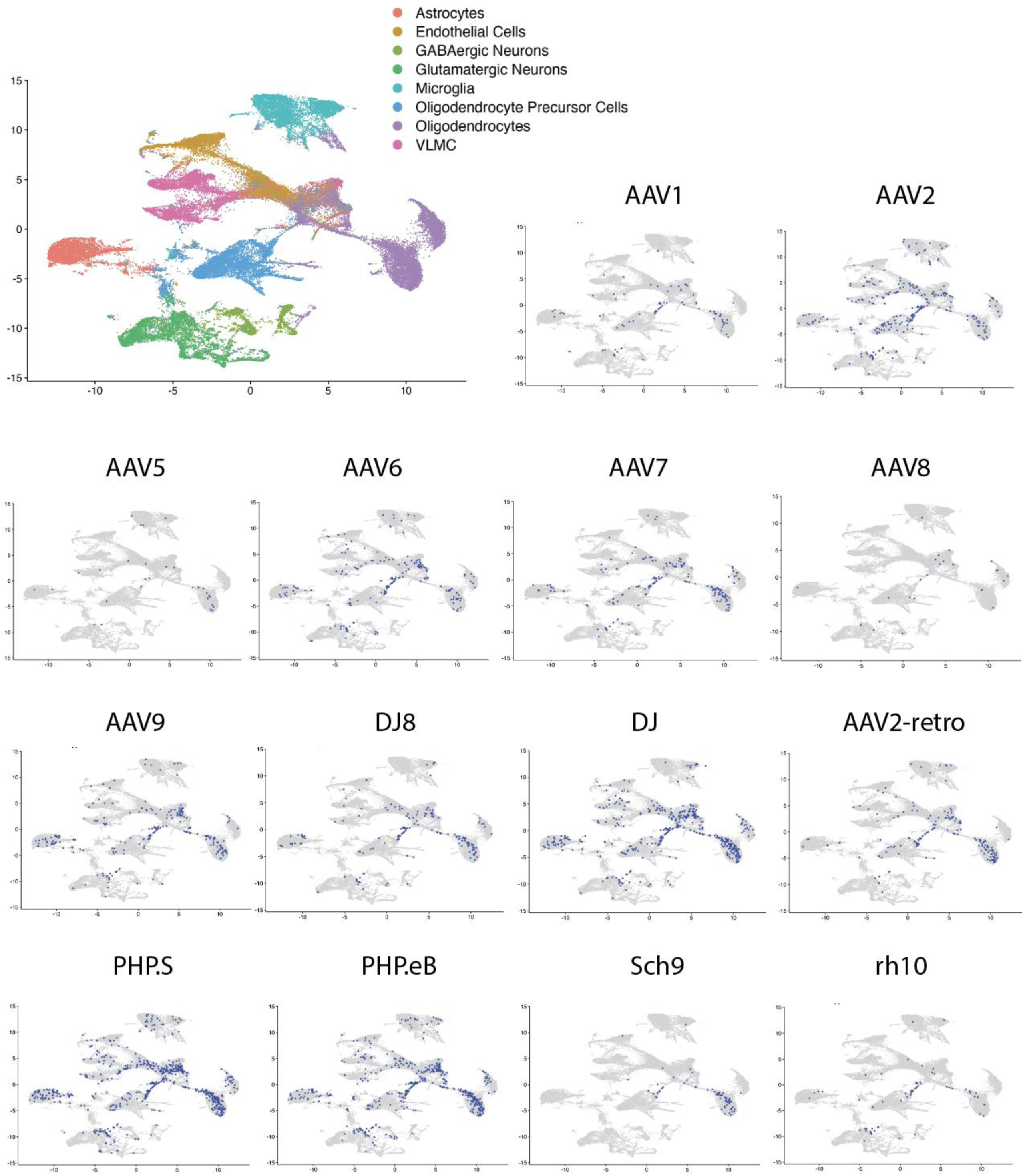
| All AAV capsid variants show broad transduction across cell types. UMAPs showing the transduction profiles of each AAV capsid variant across cell types (purple dot = cells with at least one viral transcript detected). All show transduction of numerous cell types.

**Supplemental Table 1.**
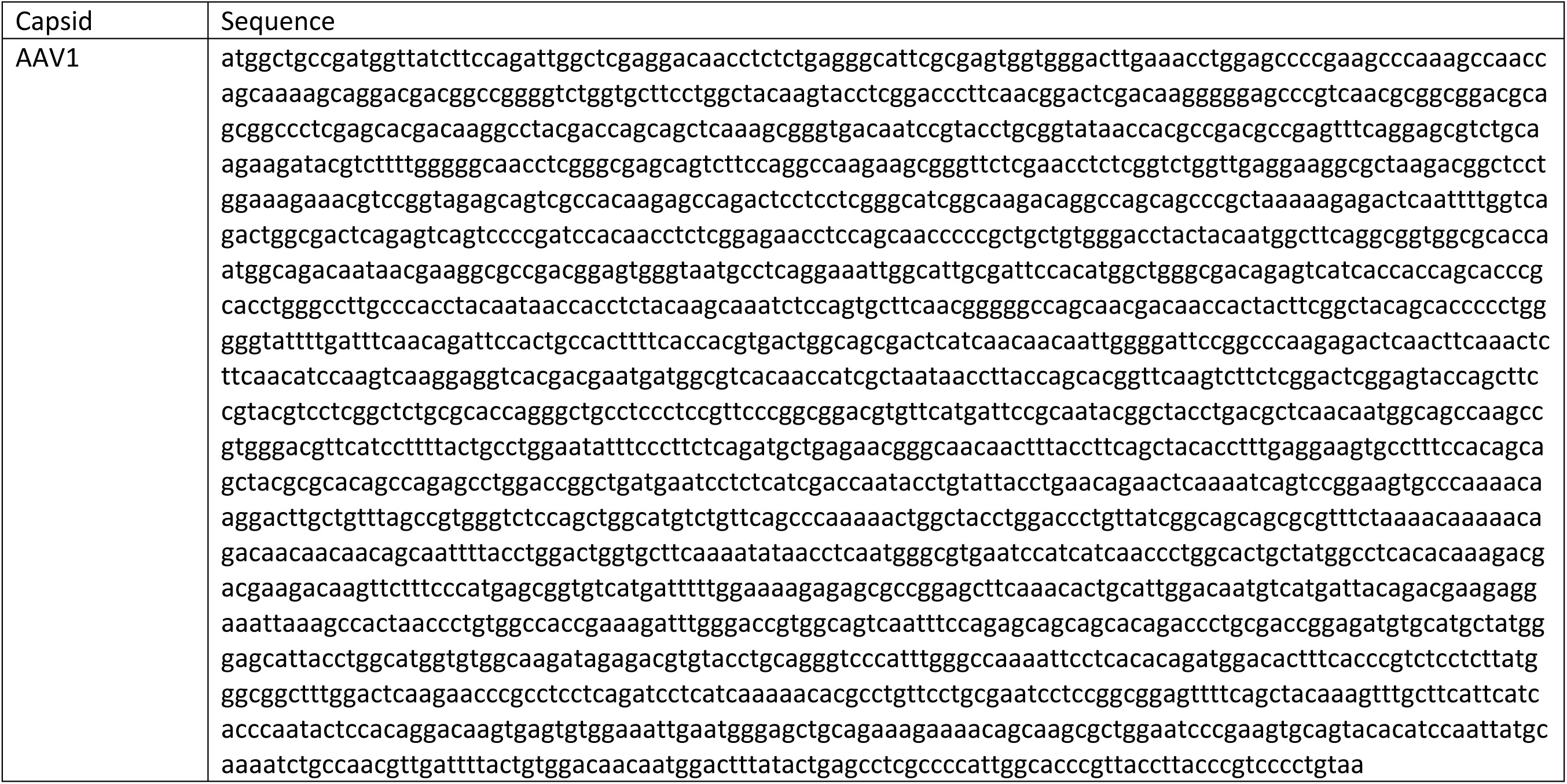
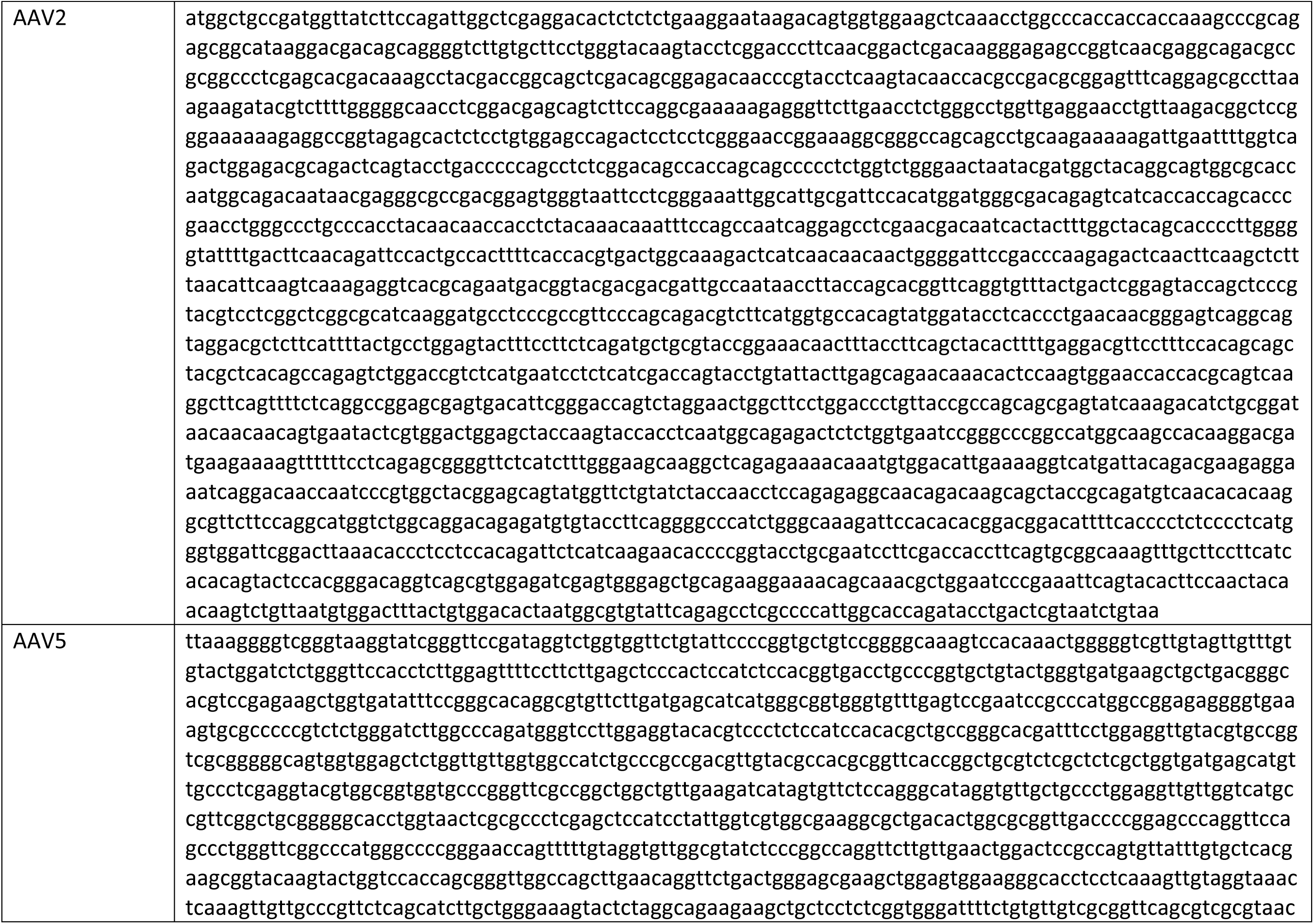
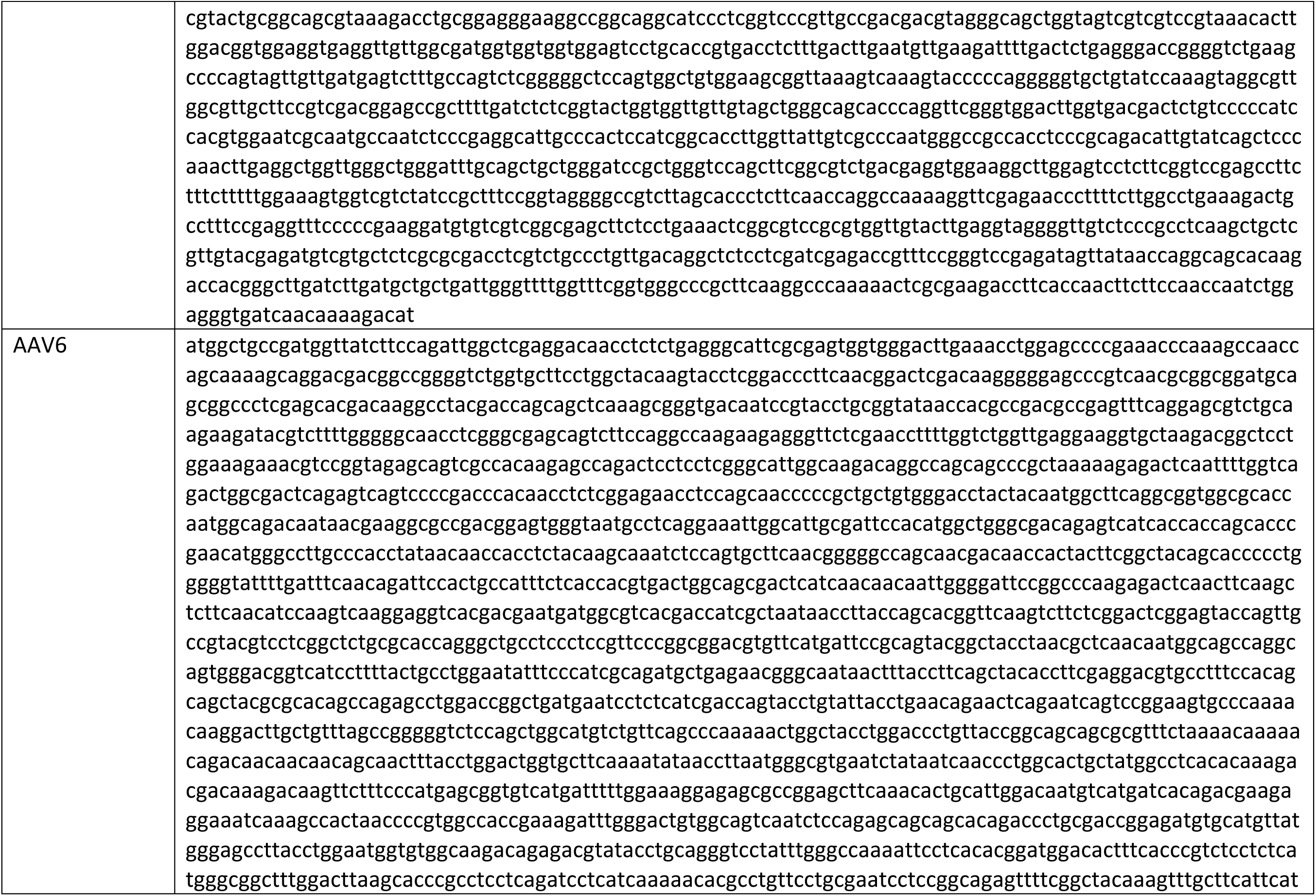
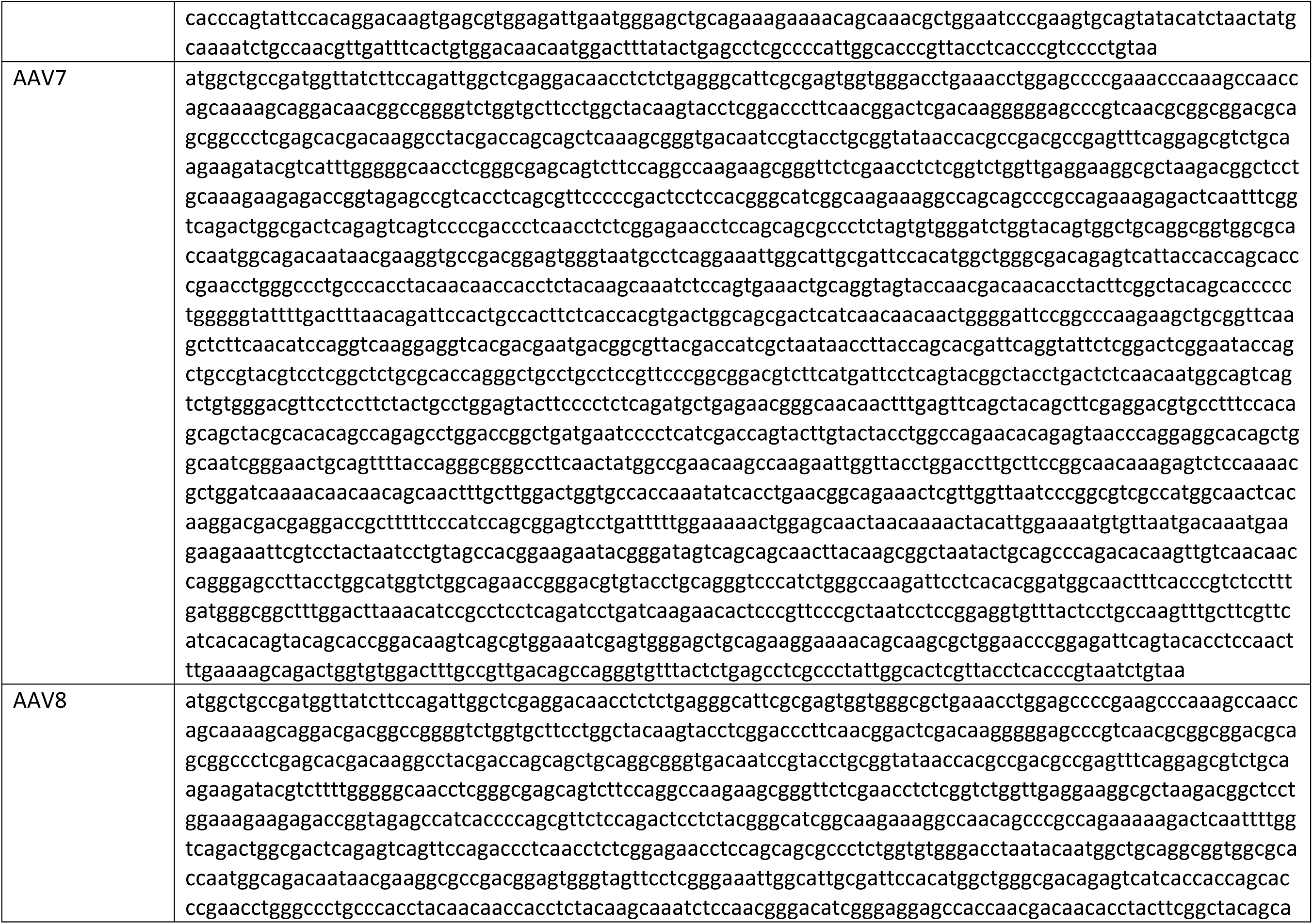
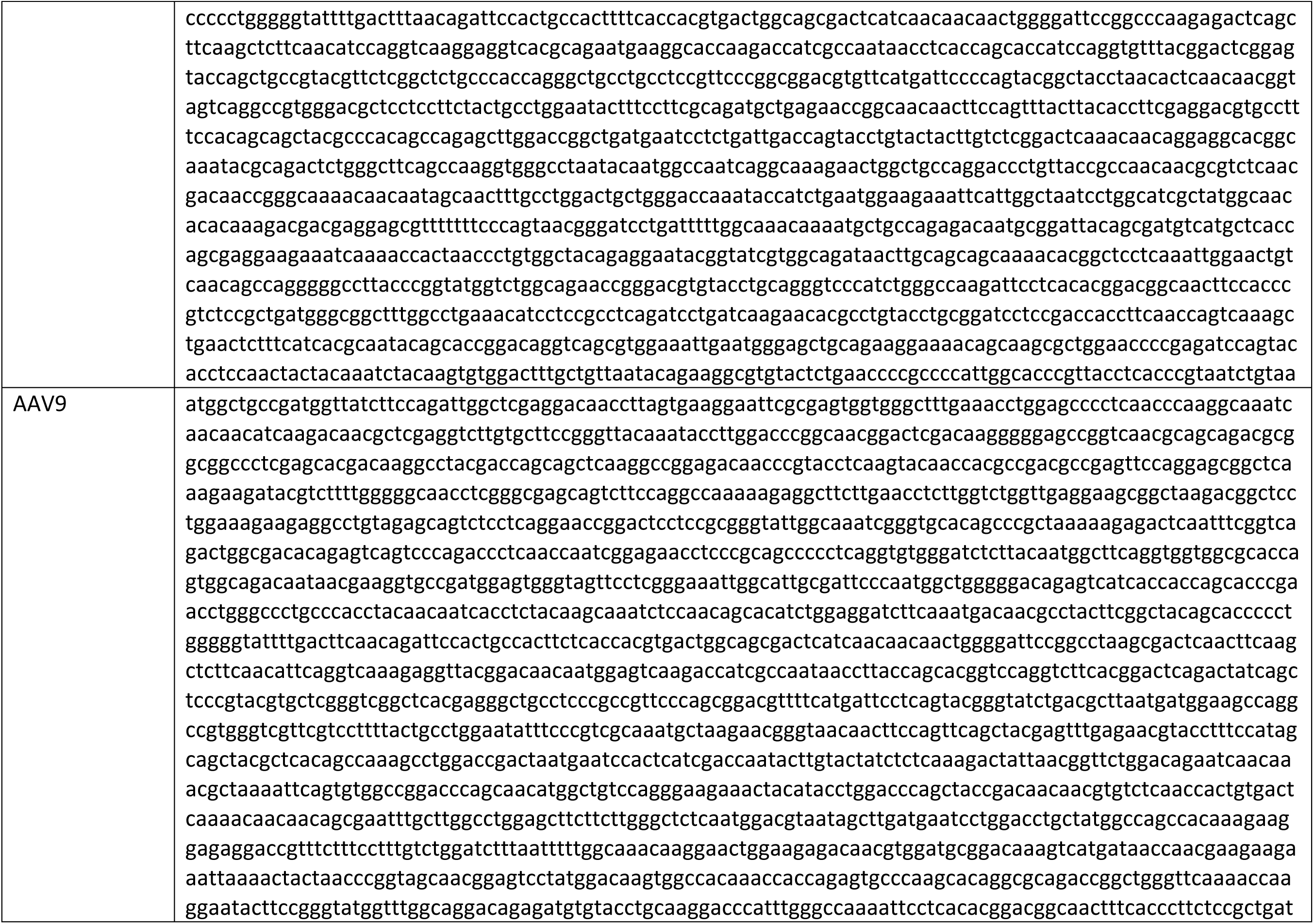
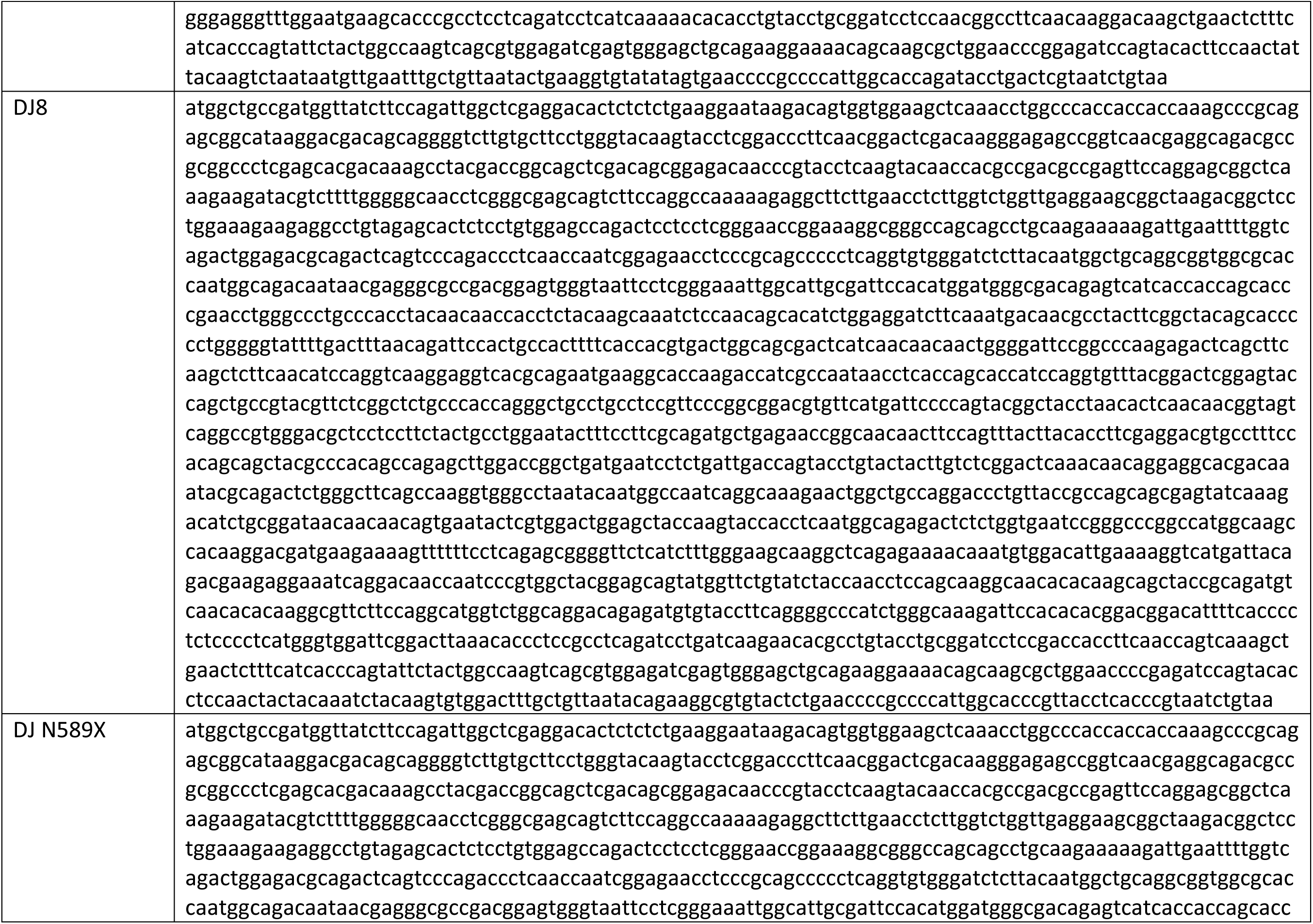
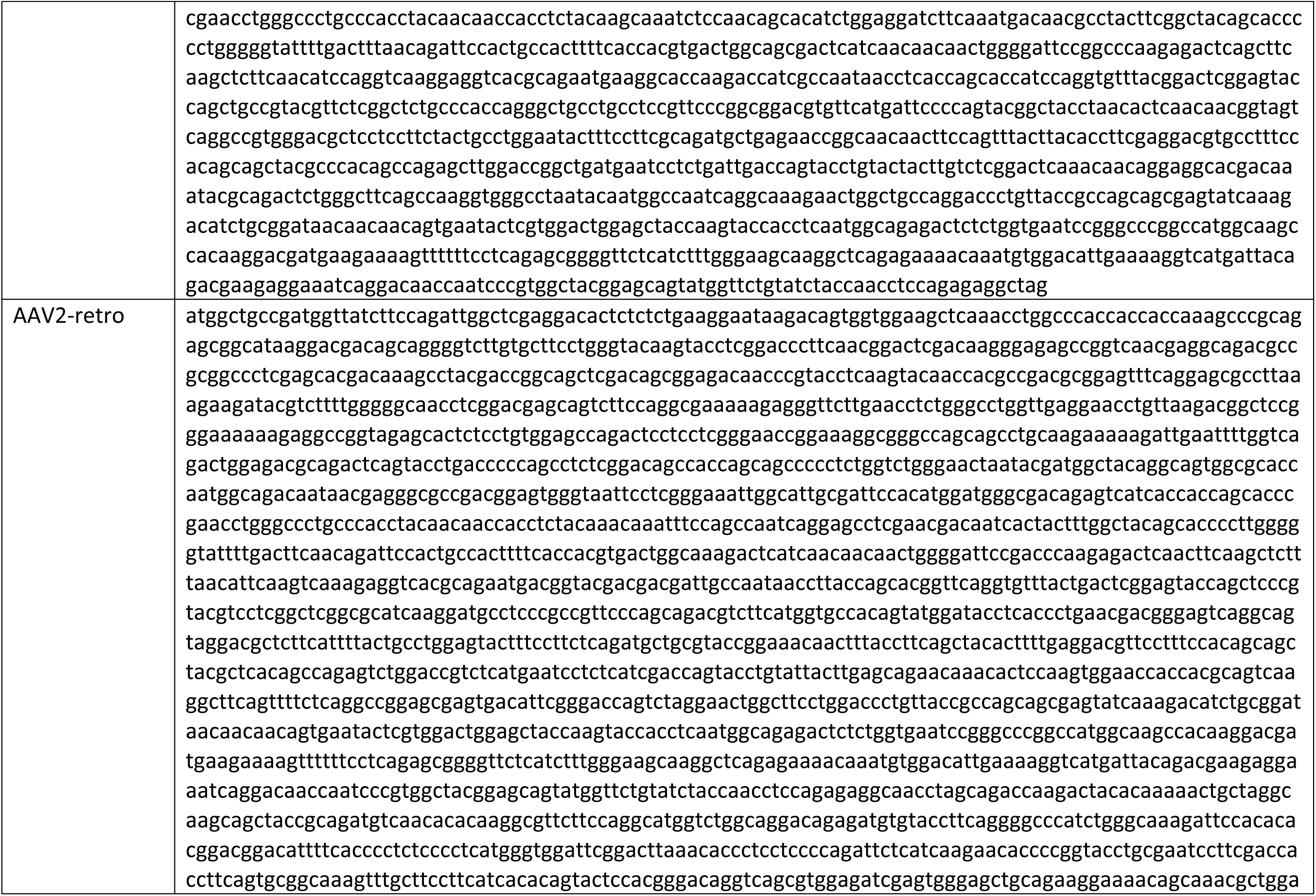
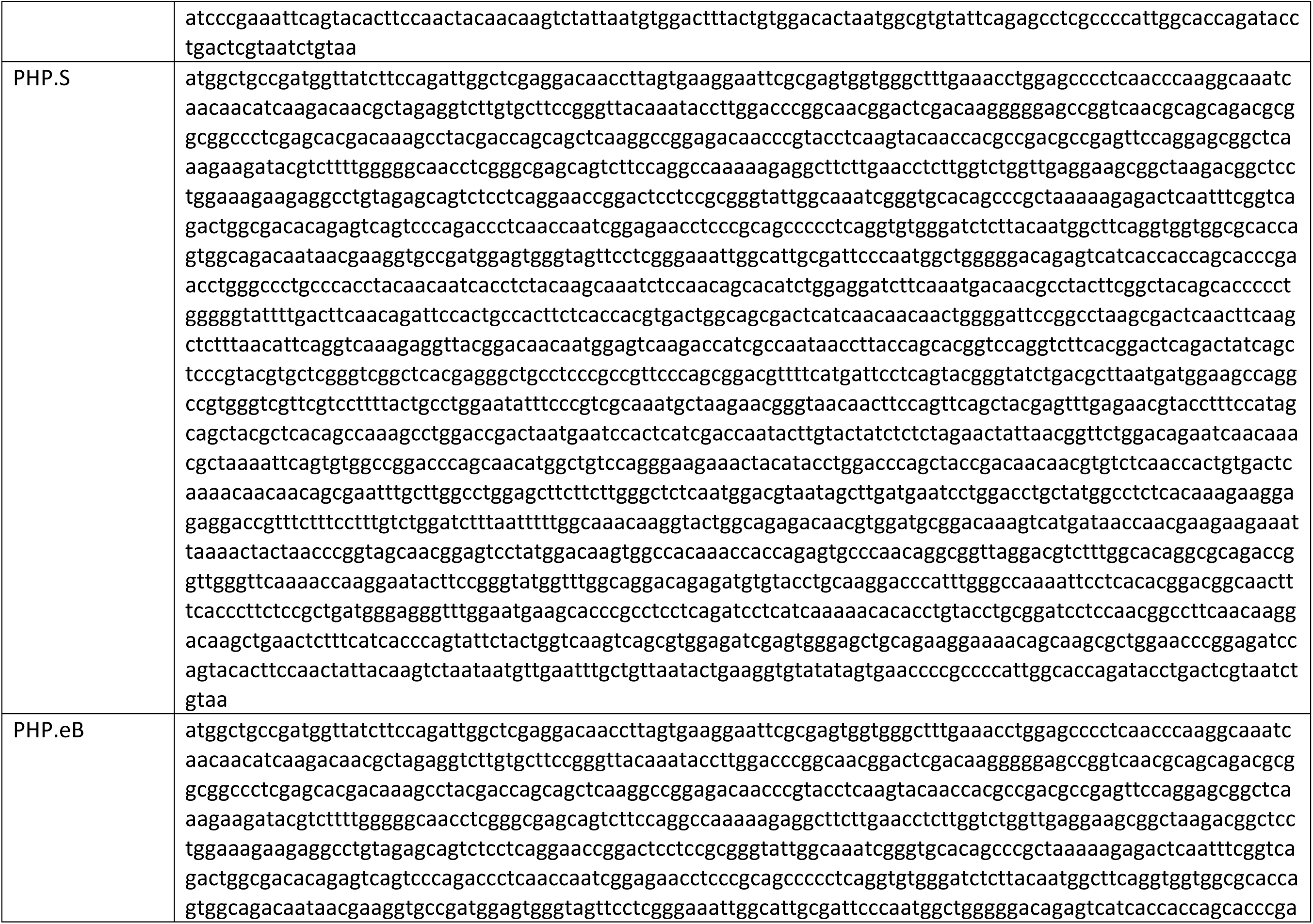
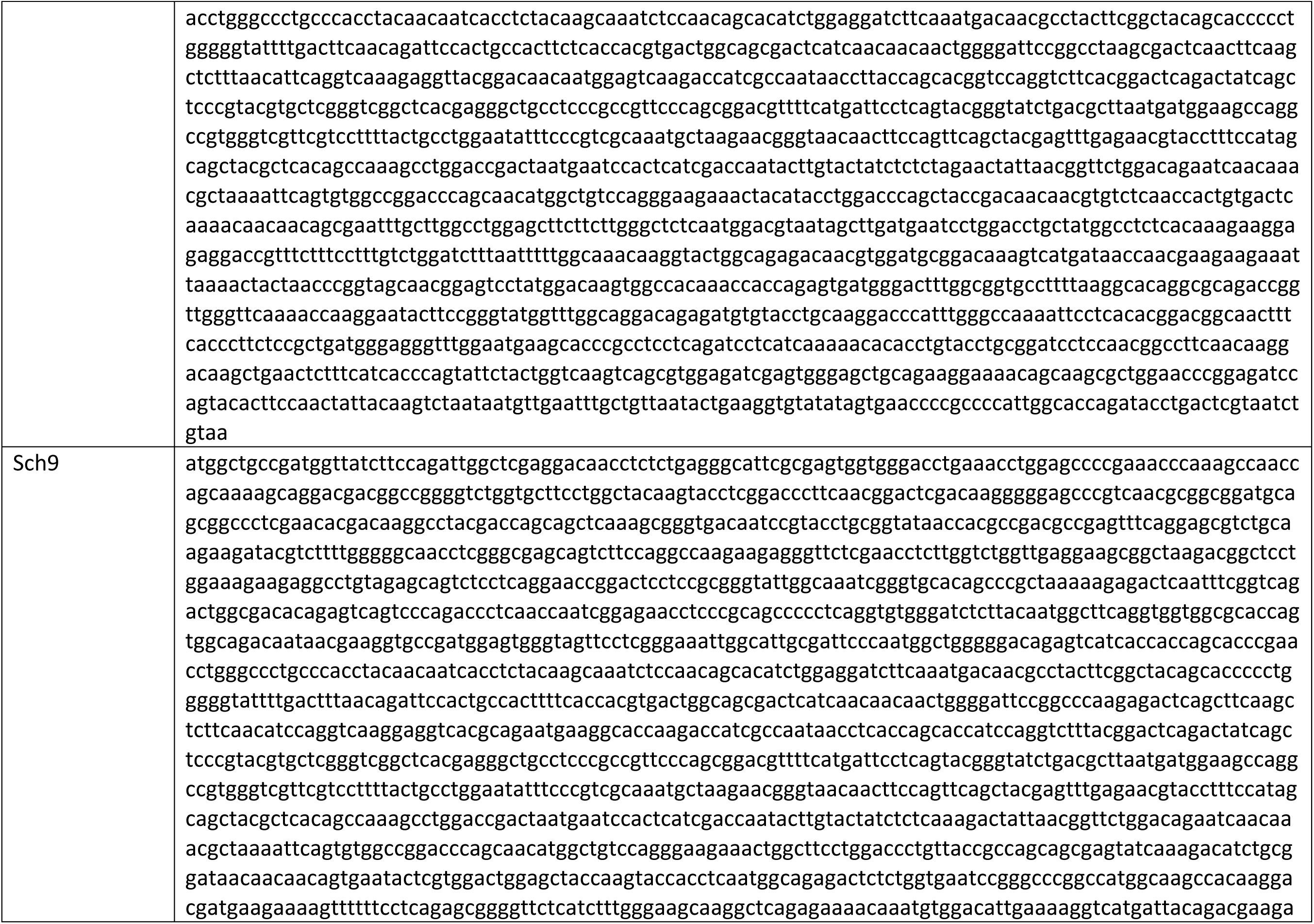
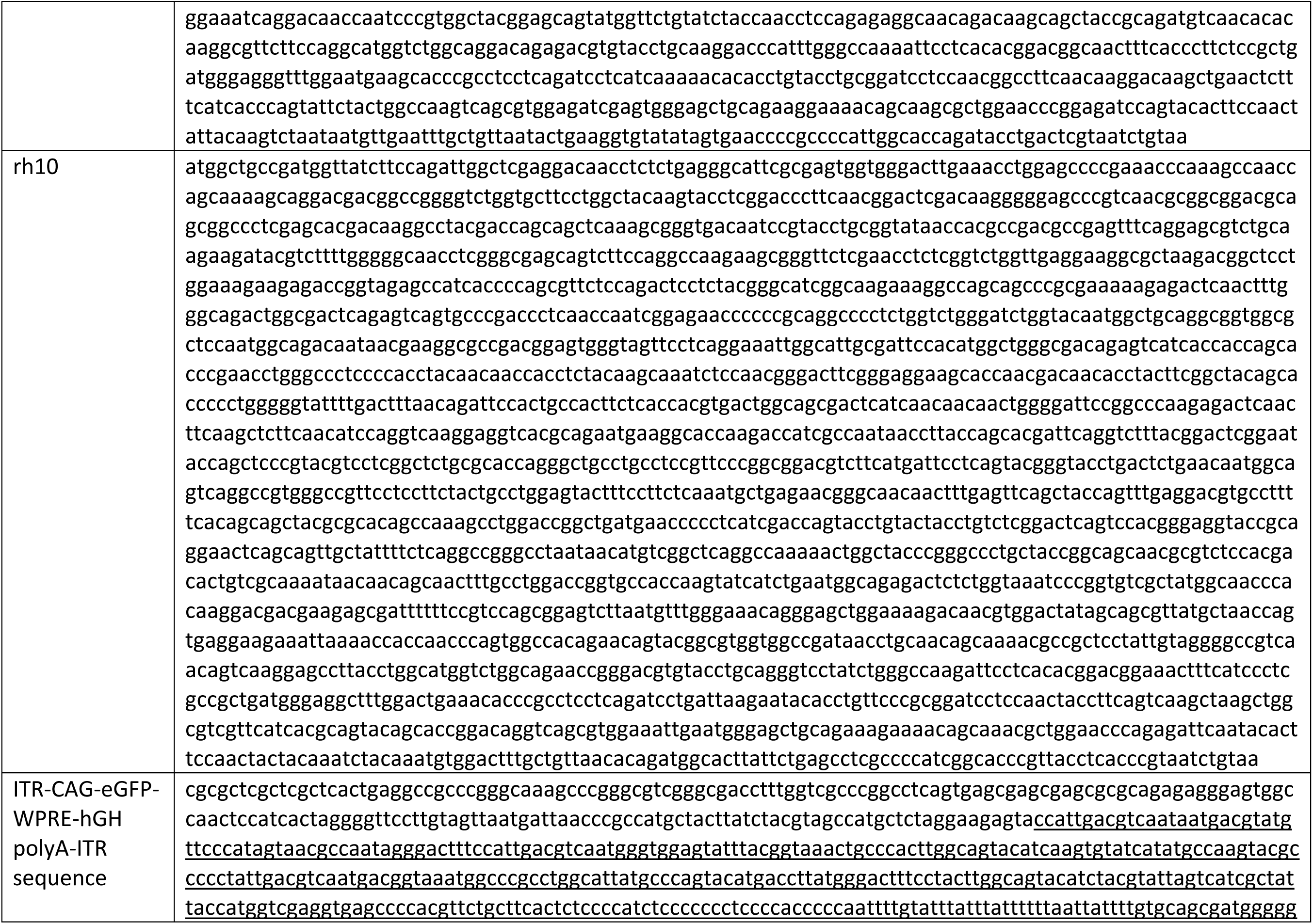
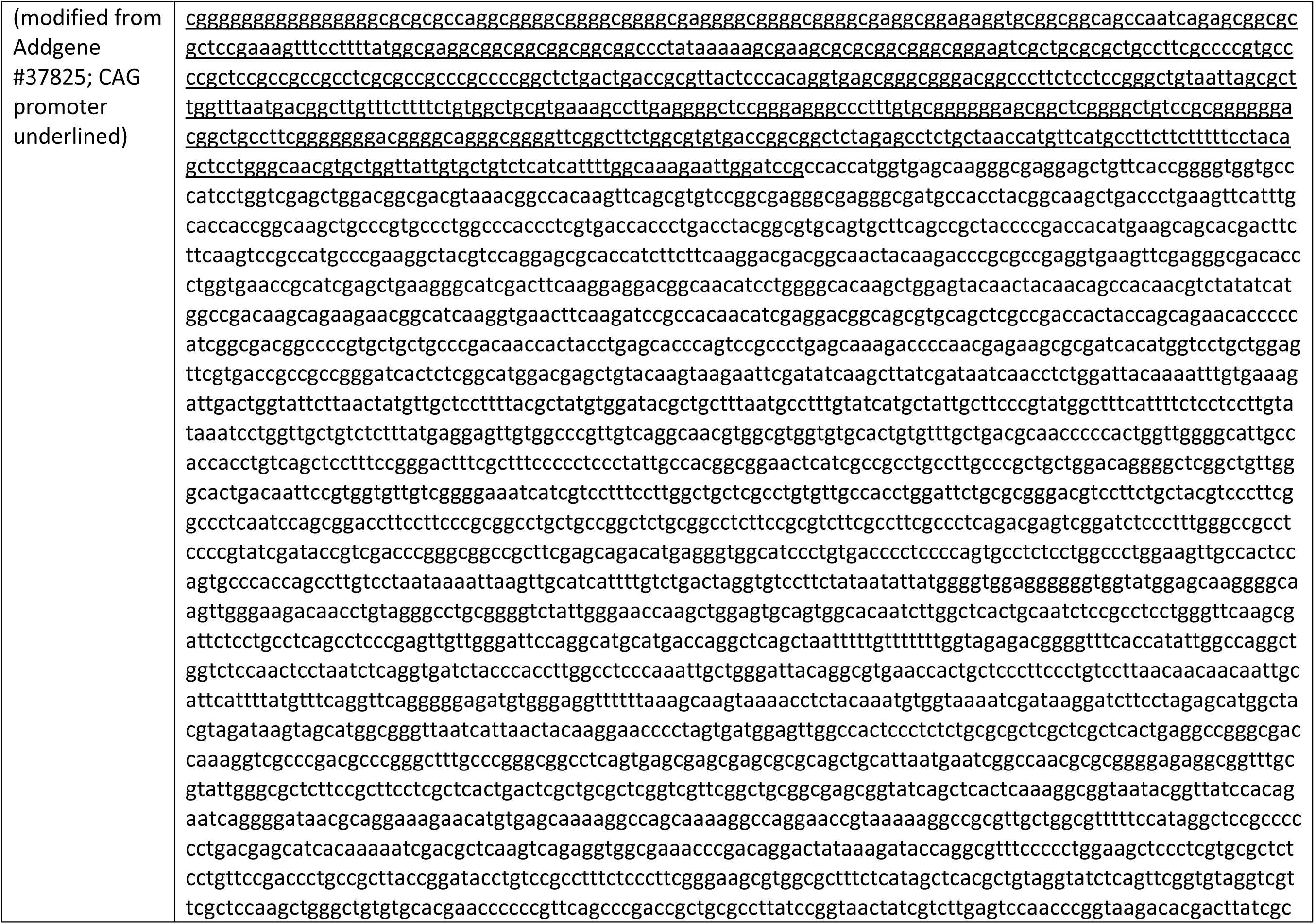
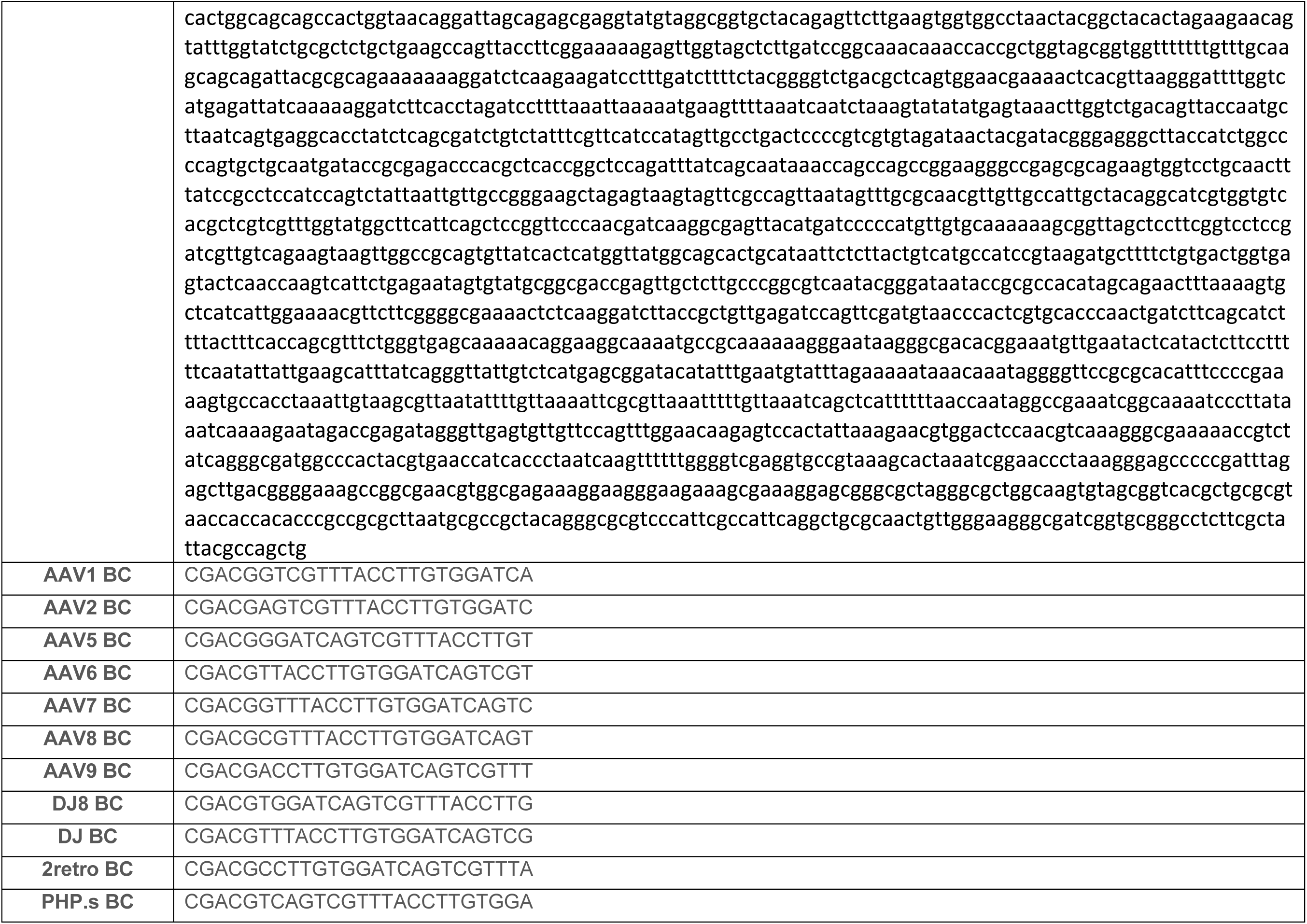
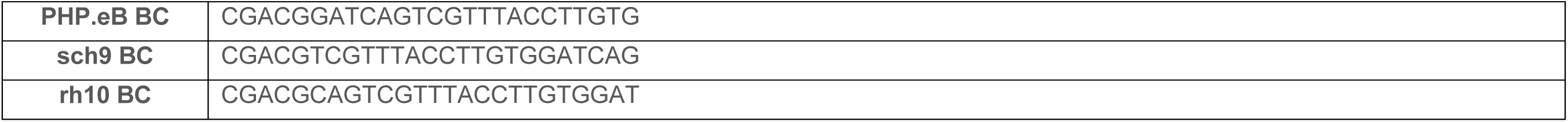
| AAV capsid, cargo, and barcode sequences used. A table with the fourteen Cap sequences used in this study, the ITR-CAG-eGFP-WPRE-hGH-ITR (modified from Addgene 37825, gift from Ed Boyden), and barcodes, corresponding to their respective capsid variants, which were placed in between the WPRE and hGH sequences at the NotI restriction site.

**Supplemental Table 2.** | Estimates of intraparenchymal trial viral dosing per mg of brain tissue. Using published estimates of diffusion spread during convection-enhanced delivery and the mass of brain tissue, we calculated the estimated range of vector genomes per mg of brain tissue dosed in intraparenchymal trials for which data could be found, assuming 1-3 µLof brain infused per µL infusate. Given that convection-enhanced delivery generally infuses an area uniformly, and passive application as we did here mainly exposes the surface of the tissue slice to virus, we think that ∼10-20% of our tissue slice is seeing the virus at its highest concentration, and so a fraction of the 10-20 mg slice (around 1-3 mg) is truly being dosed by the 2.1E9, similar to the median vg/mg in convection-enhanced trials.

| Trial | NCT | Low estimate, vector genomes per mg brain infused* | High vector genomes per mg brain infused** |
| --- | --- | --- | --- |
| NTN Phase 1a, 2a | NCT00252850 | 1.05E+09 | 3.16E+09 |
| NTN Phase 1b, 2b | NCT00252850 | 9.71E+08 | 2.91E+09 |
| VY-AADC01 phase 1 UCSF cohort 1 | NCT01973543 | 5.39E+08 | 1.62E+09 |
| VY-AADC01 phase 1 UCSF cohort 2 | NCT01973543 | 5.39E+08 | 1.62E+09 |
| VY-AADC01 phase 1 UCSF cohort 3 | NCT01973543 | 1.69E+09 | 5.07E+09 |
| GAD phase 1 2007 Lancet low dose | NCT00195143 | 4.47E+08 | 1.34E+09 |
| GAD phase 1 2007 Lancet mid dose | NCT00195143 | 1.34E+09 | 4.02E+09 |
| GAD phase 1 2007 Lancet high dose | NCT00195143 | 4.47E+09 | 1.34E+10 |
| GAD phase 2 | NCT00643890 | 3.24E+07 | 9.71E+07 |
| Tay Sachs Phase 1/2 abstract CNS 2023 | NCT04669535 | 1.06E+10 | 3.17E+10 |
| NTN Sangamo, AAV2-NRTN in PD | NCT00985517 | 2.16E+09 | 6.47E+09 |
| MPS, rh.10, SGSH | NCT03612869 | 3.24E+08 | 9.71E+08 |
| **Median** |  | **1.01E+09** | **3.03E+09** |
| Mean |  | 2.01E+09 | 6.03E+09 |

\*Low estimate of vg/mg tissue: (#vector genomes infused)/( μL of infusate \* 3 \* 1.03mg)
\*\*High estimate of vg/mg tissue: (#vector genomes infused)/( μL of infusate \* 1 \* 1.03mg)

## Notes

### Competing Interest Statement

The authors have declared no competing interest.

## References

1. Lonser RR, Akhter AS, Zabek M, Elder JB, Bankiewicz KS. Direct convective delivery of adeno-associated virus gene therapy for treatment of neurological disorders. J Neurosurg. 2020 Jul 10;134(6):1751–1763. doi: 10.3171/2020.4.JNS20701. PMID: 32915526.

2. Challis RC, Ravindra Kumar S, Chen X, Goertsen D, Coughlin GM, Hori AM, Chuapoco MR, Otis TS, Miles TF, Gradinaru V. Adeno-Associated Virus Toolkit to Target Diverse Brain Cells. Annu Rev Neurosci. 2022 Jul 8;45:447–469. doi: 10.1146/annurev-neuro-111020-100834. Epub 2022 Apr 19. PMID: 35440143.

3. Patel RV, Nanda P, Richardson RM. Neurosurgical gene therapy for central nervous system diseases. Neurotherapeutics. 2024 Aug 26;21(4):e00434. doi: 10.1016/j.neurot.2024.e00434. Epub ahead of print. PMID: 39191071.

4. Kreatsoulas D, Damante M, Cua S, Lonser RR. Adjuvant convection-enhanced delivery for the treatment of brain tumors. J Neurooncol. 2024 Jan;166(2):243–255. doi: 10.1007/s11060-023-04552-8. Epub 2024 Jan 23. PMID: 38261143; PMCID: PMC10834622.

5. Pupo A, Fernández A, Low SH, François A, Suárez-Amarán L, Samulski RJ. AAV vectors: The Rubik’s cube of human gene therapy. Mol Ther. 2022 Dec 7;30(12):3515–3541. doi: 10.1016/j.ymthe.2022.09.015. Epub 2022 Oct 5. PMID: 36203359; PMCID: PMC9734031.

6. Ravindra Kumar S, Miles TF, Chen X, Brown D, Dobreva T, Huang Q, Ding X, Luo Y, Einarsson PH, Greenbaum A, Jang MJ, Deverman BE, Gradinaru V. Multiplexed Cre-dependent selection yields systemic AAVs for targeting distinct brain cell types. Nat Methods. 2020 May;17(5):541–550. doi: 10.1038/s41592-020-0799-7. Epub 2020 Apr 20. PMID: 32313222; PMCID: PMC7219404.

7. Ogden PJ, Kelsic ED, Sinai S, Church GM. Comprehensive AAV capsid fitness landscape reveals a viral gene and enables machine-guided design. Science. 2019 Nov 29;366(6469):1139-1143. doi: 10.1126/science.aaw2900. PMID: 31780559; PMCID: PMC7197022.

8. Pekrun K, De Alencastro G, Luo QJ, Liu J, Kim Y, Nygaard S, Galivo F, Zhang F, Song R, Tiffany MR, Xu J, Hebrok M, Grompe M, Kay MA. Using a barcoded AAV capsid library to select for clinically relevant gene therapy vectors. JCI Insight. 2019 Nov 14;4(22):e131610. doi: 10.1172/jci.insight.131610. PMID: 31723052; PMCID: PMC6948855.

9. Li C, Samulski RJ. Engineering adeno-associated virus vectors for gene therapy. Nat Rev Genet. 2020 Apr;21(4):255–272. doi: 10.1038/s41576-019-0205-4. Epub 2020 Feb 10. PMID: 32042148.

10. Bartus RT, Herzog CD, Chu Y, Wilson A, Brown L, Siffert J, Johnson EM Jr, Olanow CW, Mufson EJ, Kordower JH. Bioactivity of AAV2-neurturin gene therapy (CERE-120): differences between Parkinson’s disease and nonhuman primate brains. Mov Disord. 2011 Jan;26(1):27–36. doi: 10.1002/mds.23442. Epub 2010 Nov 18. PMID: 21322017; PMCID: PMC6333467.

11. Chu Y, Bartus RT, Manfredsson FP, Olanow CW, Kordower JH. Long-term post-mortem studies following neurturin gene therapy in patients with advanced Parkinson’s disease. Brain. 2020 Mar 1;143(3):960–975. doi: 10.1093/brain/awaa020. PMID: 32203581; PMCID: PMC7089653.

12. Castle MJ, Baltanás FC, Kovacs I, Nagahara AH, Barba D, Tuszynski MH. Postmortem Analysis in a Clinical Trial of AAV2-NGF Gene Therapy for Alzheimer’s Disease Identifies a Need for Improved Vector Delivery. Hum Gene Ther. 2020 Apr;31(7-8):415–422. doi: 10.1089/hum.2019.367. Epub 2020 Mar 30. PMID: 32126838; PMCID: PMC7194314.

13. Chaichana KL, Capilla-Gonzalez V, Gonzalez-Perez O, Pradilla G, Han J, Olivi A, Brem H, Garcia-Verdugo JM, Quiñones-Hinojosa A. Preservation of glial cytoarchitecture from ex vivo human tumor and non-tumor cerebral cortical explants: A human model to study neurological diseases. J Neurosci Methods. 2007 Aug 30;164(2):261–70. doi: 10.1016/j.jneumeth.2007.05.008. Epub 2007 May 16. PMID: 17580092; PMCID: PMC2744592.

14. Park TI, Smyth LCD, Aalderink M, Woolf ZR, Rustenhoven J, Lee K, Jansson D, Smith A, Feng S, Correia J, Heppner P, Schweder P, Mee E, Dragunow M. Routine culture and study of adult human brain cells from neurosurgical specimens. Nat Protoc. 2022 Feb;17(2):190–221. doi: 10.1038/s41596-021-00637-8. Epub 2022 Jan 12. PMID: 35022619.

15. Ting JT, Lee BR, Chong P, Soler-Llavina G, Cobbs C, Koch C, Zeng H, Lein E. Preparation of Acute Brain Slices Using an Optimized N-Methyl-D-glucamine Protective Recovery Method. J Vis Exp. 2018 Feb 26;(132):53825. doi: 10.3791/53825. PMID: 29553547; PMCID: PMC5931343.

16. Ting JT, Kalmbach B, Chong P, de Frates R, Keene CD, Gwinn RP, Cobbs C, Ko AL, Ojemann JG, Ellenbogen RG, Koch C, Lein E. A robust ex vivo experimental platform for molecular-genetic dissection of adult human neocortical cell types and circuits. Sci Rep. 2018 May 30;8(1):8407. doi: 10.1038/s41598-018-26803-9. PMID: 29849137; PMCID: PMC5976666.

17. Schwarz N, Hedrich UBS, Schwarz H, P A H, Dammeier N, Auffenberg E, Bedogni F, Honegger JB, Lerche H, Wuttke TV, Koch H. Human Cerebrospinal fluid promotes long-term neuronal viability and network function in human neocortical organotypic brain slice cultures. Sci Rep. 2017 Sep 25;7(1):12249. doi: 10.1038/s41598-017-12527-9. PMID: 28947761; PMCID: PMC5613008.

18. Bak A, Koch H, van Loo KMJ, Schmied K, Gittel B, Weber Y, Ort J, Schwarz N, Tauber SC, Wuttke TV, Delev D. Human organotypic brain slice cultures: a detailed and improved protocol for preparation and long-term maintenance. J Neurosci Methods. 2024 Apr;404:110055. doi: 10.1016/j.jneumeth.2023.110055. Epub 2024 Jan 5. PMID: 38184112.

19. Allen Institute for Brain Science. Cell Types Database: RNA-Seq Data. [Internet]. Seattle (WA): Allen Institute for Brain Science; c2024 [cited 2024 Sep 11]. Available from: https://celltypes.brain-map.org/rnaseq/Human-MTG-10x_SEA-AD?selectedVisualization=Heatmap&colorByFeature=Cell+Type&colorByFeatureValue=GAD1

20. Aschauer DF, Kreuz S, Rumpel S. Analysis of transduction efficiency, tropism and axonal transport of AAV serotypes 1, 2, 5, 6, 8 and 9 in the mouse brain. PLoS One. 2013 Sep 27;8(9):e76310. doi: 10.1371/journal.pone.0076310. PMID: 24086725; PMCID: PMC3785459.

21. He T, Itano MS, Earley LF, Hall NE, Riddick N, Samulski RJ, Li C. The Influence of Murine Genetic Background in Adeno-Associated Virus Transduction of the Mouse Brain. Hum Gene Ther Clin Dev. 2019 Dec;30(4):169–181. doi: 10.1089/humc.2019.030. PMID: 31749390; PMCID: PMC6919261.

22. Schwanhäusser B, Busse D, Li N, Dittmar G, Schuchhardt J, Wolf J, Chen W, Selbach M. Global quantification of mammalian gene expression control. Nature. 2011 May 19;473(7347):337–42. doi: 10.1038/nature10098. Erratum in: Nature. 2013 Mar 07;495(7439):126-7. doi: 10.1038/nature11848. PMID: 21593866.

23. Gonzalez-Sandoval A, Pekrun K, Tsuji S, Zhang F, Hung KL, Chang HY, Kay MA. The AAV capsid can influence the epigenetic marking of rAAV delivered episomal genomes in a species dependent manner. Nat Commun. 2023 Apr 28;14(1):2448. doi: 10.1038/s41467-023-38106-3. PMID: 37117181; PMCID: PMC10147666.

24. Loeb EJ, Havlik PL, Elmore ZC, Rosales A, Fergione SM, Gonzalez TJ, Smith TJ, Benkert AR, Fiflis DN, Asokan A. Capsid-mediated control of adeno-associated viral transcription determines host range. Cell Rep. 2024 Mar 26;43(3):113902. doi: 10.1016/j.celrep.2024.113902. Epub 2024 Mar 2. PMID: 38431840; PMCID: PMC11150003.

25. Harkins AL, Ambegaokar PP, Keeler AM. Immune responses to central nervous system directed adeno-associated virus gene therapy: Does direct CNS delivery make a difference? Neurotherapeutics. 2024 Aug 23;21(4):e00435. doi: 10.1016/j.neurot.2024.e00435. Epub ahead of print. PMID: 39180957.

26. Schwarz N, Uysal B, Welzer M, Bahr JC, Layer N, Löffler H, Stanaitis K, Pa H, Weber YG, Hedrich UB, Honegger JB, Skodras A, Becker AJ, Wuttke TV, Koch H. Long-term adult human brain slice cultures as a model system to study human CNS circuitry and disease. Elife. 2019 Sep 9;8:e48417. doi: 10.7554/eLife.48417. PMID: 31498083

27. Scannell JW, Bosley J. When Quality Beats Quantity: Decision Theory, Drug Discovery, and the Reproducibility Crisis. PLoS One. 2016 Feb 10;11(2):e0147215. doi: 10.1371/journal.pone.0147215. PMID: 26863229; PMCID: PMC4749240.

28. Zhu D, Brookes DH, Busia A, Carneiro A, Fannjiang C, Popova G, Shin D, Donohue KC, Lin LF, Miller ZM, Williams ER, Chang EF, Nowakowski TJ, Listgarten J, Schaffer DV. Optimal trade-off control in machine learning-based library design, with application to adeno-associated virus (AAV) for gene therapy. Sci Adv. 2024 Jan 26;10(4):eadj3786. doi: 10.1126/sciadv.adj3786. Epub 2024 Jan 24. PMID: 38266077

